# Remote force modulation of the T-cell receptor reveals an NFAT-threshold for CD4^+^ T cell activation

**DOI:** 10.1101/2024.10.20.619286

**Authors:** Joseph Clarke, Jeremy Pike, David Bending, Dylan Owen, David C. Wraith, Alicia J El Haj

**Author notes:** MRC Translational Immune Discovery Unit, Radcliffe Department of Medicine, John Radcliffe Hospital, University of Oxford, Oxford, OX3 9DS, United Kingdom. Corresponding authors: Joseph Clarke;, Alicia J El Haj.

## Abstract

Mechano-modulation of cell surface proteins to influence cell activation has been shown as a promising new advanced therapy for regenerative medicine applications. These strategies rely on the manipulation of mechanosensitive cell surface receptors to initiate intracellular signal transduction. The cell surface receptor of T lymphocytes (TCR), which recognises peptide-MHC molecules central to driving the adaptive immune response, has recently been suggested to be mechano-responsive. Despite this advance, little is known as to whether the TCR can be mechanically modulated to achieve TCR signalling and subsequent T cell activation, and whether these characteristics can be exploited for immunotherapies. Here, we describe a magnetic particle-based platform for mechanical modulation of the TCR and outline how this platform can be utilised to achieve CD4^+^ T cell activation. We demonstrate that mechanical manipulation of the TCR induces cell surface clustering of the TCR and downstream TCR signalling, leading to eventual TCR downregulation and T cell activation. We investigate the temporal relationship between mechanical modulation of the TCR and subsequent T cell activation, hereby identifying that accumulation of signalling events within the NFAT-pathway is required to reach the threshold required for CD4^+^ T cell activation, outlining an axis which controls the CD4^+^ T cell response to external mechanical cues. These findings identify how CD4^+^ T cells can modulate their function in response to such cues, whilst also outlining a remote-magnetic particle-based platform that may be used for the control of T cell responses.

## Introduction

T cell subtypes form a critical part of the adaptive immune response. Antigen recognition by T lymphocytes, achieved by recognition of peptide-MHC (pMHC) molecules on the surface of antigen presenting cells (APCs) by the T cell receptor (TCR), forms a critical part of this response. Multiple models have been proposed to explain how TCR triggering can occur, including TCR aggregation-based models^1–4^, kinetic segregation (KS) models^5–7^ and kinetic proof-reading (KPR) models^8–11^, and each provide plausible mechanisms by which TCR triggering can occur. Briefly, these models indicate how arrangement of proteins in, or immediately proximal to, the T cell membrane can influence TCR triggering and T cell activation. The KS model postulates that segregation of TCR-CD3 complexes from transmembrane phosphatases, resulting in size-based exclusion from close contact regions between T cell and APC on account of the large ectodomains of these phosphatases, shifts the kinase-phosphatase activity balance in the favour of CD3 phosphorylation allowing for TCR signalling ^5–7^. Similarly, aggregation-based models propose that TCR-CD3 aggregation in the cell membrane can increase local proximity of these molecules with activating kinases such as Lck (reported to be both membrane and co-receptor associated), and hence also induce TCR triggering and signalling ^1–4^. Finally, KPR models centre around the notion that only TCR-pMHC interactions of a suitable affinity will allow for bond life-times of a sufficient time frame (in the order of seconds) for these initiating events to occur, and that weaker interactions that do not survive this frame will dissociate before signalling can occur ^8–11^.

However, recently it has been proposed that the TCR may function as a mechanosensor, whereby force application to the TCR-CD3 complex may directly induce TCR signalling. Recent studies utilising both optical tweezers and atomic force microscopy to apply force to the TCR-CD3 complex have shown evidence of force induced calcium signalling downstream of the TCR^12–14^. Importantly, bonds between TCR and agonistic pMHC have been shown to form catch bonds, whereby bond lifetimes are stabilised and hence prolonged under force, whilst TCR interactions within non-specific pMHC are shown to form slip bonds and instead dissociate under force, presumably before phosphorylation events postulated by the KPR model can take place^15–17^. These reports hence implicate how force application to the TCR-pMHC interaction can play a direct role in antigen discrimination. Despite these advances, it is poorly established how force application to the TCR-CD3 complex can influence signalling and T cell activation beyond the level of calcium signalling, an important point to establish since not all TCR-mediated calcium fluxes will result in robust T cell activation. Additionally, responses to mechanical changes in the environment by T cells has been shown to be important during the course of infection, fibrosis, and cancer, outlining that altered mechanical environments within which T cell activation takes place can shape the T cell response^18^.

In line with this, remote mechanical modulation of cell surface receptors has been shown to be a promising therapeutic approach in regenerative medicine and other neuro-diseases^19–23^. Here, we use this novel approach to demonstrate that the TCR can be activated with a remote magnetic-particle based method for force application to the TCR-CD3 complex. Utilising superparamagnetic nanoparticles (MNPs) functionalised with anti-CD3 antibodies exposed to external, remote dynamic magnetic field gradients, we show that force application to the TCR is able to specifically induce CD4^+^ T cell signalling. We outline that force application promotes cell surface TCR re-organisation, resulting in TCR clustering, TCR signalling and ultimately results in the removal of the TCR from the cell surface. Importantly, we define that force application for one hour is required for this response and drives NFAT-signal accumulation required to reach the threshold for CD4^+^ T cell activation, identifying signal accumulation along the NFAT pathway as a potential mechanism controlling the CD4^+^ T cell response in external mechanical cues.

## Results

### Properties of functionalised MNPs used to induce TCR signalling

We based our methodology for force application to the TCR on the previously published work utilising superparamagnetic nanoparticles (hence forth called MNPs) to achieve force application to various ion channels and mechano-receptors^19,24–26^. Functionalised MNPs, either 250nm or 1µm in diameter, targeting the CD3ε chain of the TCR-CD3 complex were utilised throughout this investigation. To achieve force application to the TCR, we made use of an external oscillating magnetic field (termed MICA, <u>M</u>agnetic Ion <u>C</u>hannel <u>A</u>ctivation, MICA Biosystems Ltd), to deliver dynamic remote magnetic field gradients^24,26^. Briefly, MICA functions as a computer programmable oscillating magnetic array, that in the ‘ON’ position transfers a magnetic force to functionalised MNPs, whilst in the ‘OFF’ position, the array is moved at a position far enough from the cell-MNP interactions such that no force is transferred. In the ‘ON’ position, the magnetic array was positioned approximately 5mm away from the cell-MNP interactions and exerted a magnetic field of approximately 91mT with a gradient of 11.1Tm^-1^. Use of MICA for external force application is illustrated in figure 1, whilst approximate forces experienced by either particle, determined as per previous reports^27,28^, are displayed in table 1.

**Figure 1.**
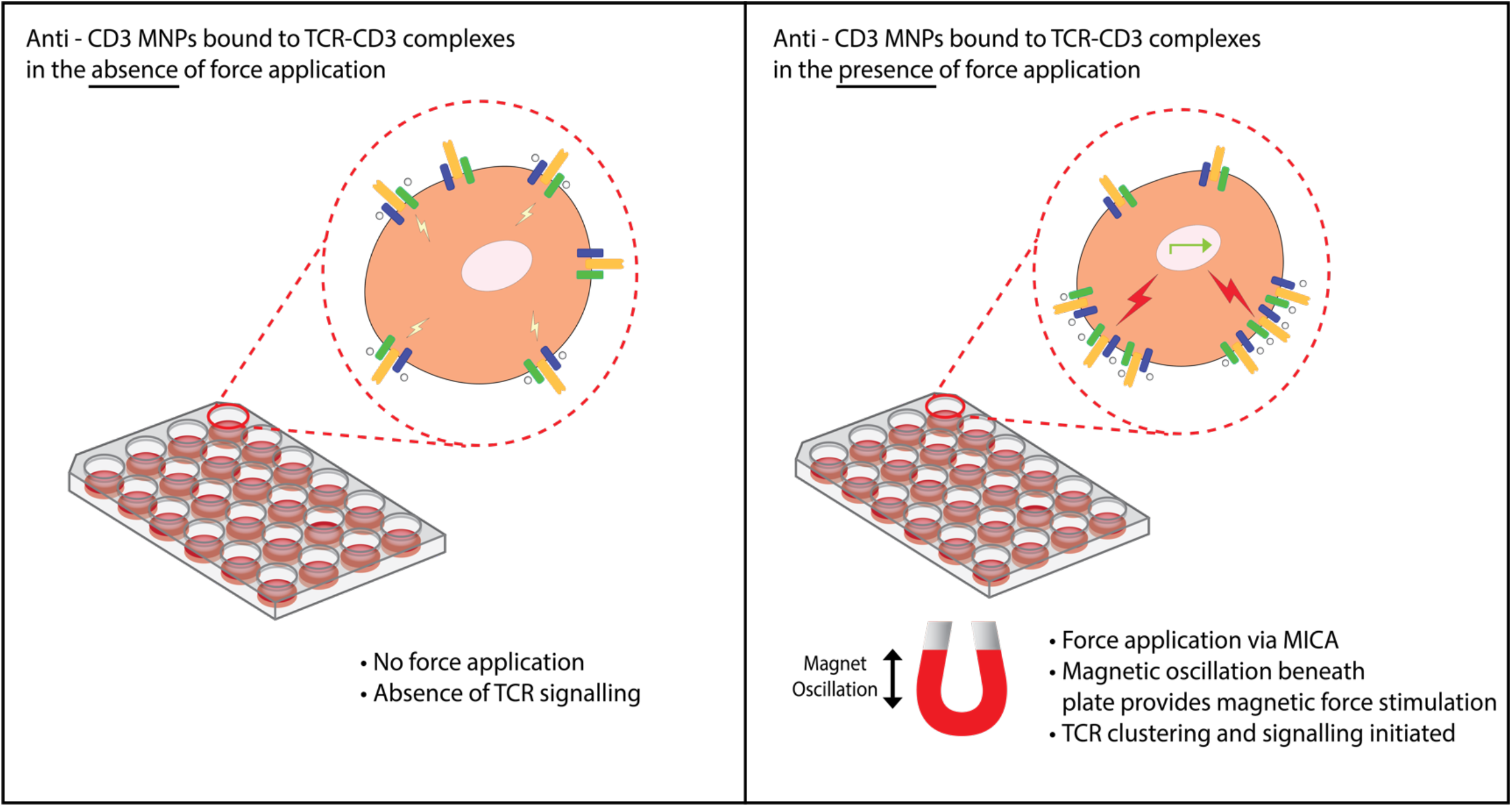
Graphical representation of force application with MICA. The MICA apparatus of external remote magnetic force application consists of an external oscillating magnet, that oscillates with a frequency of 1Hz underneath a tissue culture plate. Force was applied to CD4^+^ T cells bound to anti-CD3 or anti-CD3/CD28 MNPs for time frames indicated throughout, before measurement of biological response and T cell activation.

**Table 1.** Magnetic field properties and approximate force magnitude transferred to functionalised magnetic nanoparticles.

| Particle Size<br>(diameter) | Magnetic Field<br>Strength (mT) | Magnetic Field<br>Gradient (Tm<sup>-1</sup>) | Approximate force<br>(pN) |
| --- | --- | --- | --- |
| 250nm | 91 | 11.1 | 1.053 |
| 1µm | 91 | 11.1 | 16.848 |

To confirm functionalisation with anti-CD3ε antibodies (clonotype 145-2C11) and ensure batch-to-batch consistency in level of bound surface antibody, functionalised particles were counterstained with FITC conjugated anti-Armenian hamster IgG antibodies and analysed by flow cytometry to confirm the presence of the 145-2C11 antibody on the surface of the MNPs. Positive FITC signals identified the presence of anti-CD3ε antibodies on the surface of the particles (supplementary figures 1 and 2). This approach was also utilised to validate a level of antibody immobilised to the MNP surface sufficient to induce TCR signalling to the level where effect of magnetic force could be visualised (supplementary figure 1). Here, MNPs were functionalised as described, using either 0.1μg, 1μg, or 10μg total anti-CD3ε in the functionalisation reaction. Use of these particles to stimulate naïve murine CD4^+^ T cells showed that use of 1μg of total antibody in the functionalisation reaction was sufficient to induce TCR signalling to levels allowing for effects of magnetic force application to be observed. Decreasing this 10-fold to 0.1μg failed to induce any detectable TCR signalling (supplementary figure 1C), whilst increasing 10-fold to 10μg appeared to saturate levels of observed TCR signalling (supplementary figure 1A). Therefore, MNPs were subsequently functionalised using 1μg of total antibody in the functionalisation reaction throughout. Assuming 100% efficiency of binding of antibody to particle surface, approximations of number of antibodies per particle (for particles with diameters of 250nm or 1μm) are shown in table 2.

**Table 2.** Antibodies per surface area unit calculations for 250nm and 1µm functionalised magnetic nanoparticles.

| Particle | Amount<br>of<br>antibody<br>added | Total number<br>of antibody<br>molecules<br>added | Amount of<br>particles<br>functionalised | Stock particle<br>concentration | Antibodies<br>per<br>particle | Particle<br>surface<br>area | Antibodies<br>per<br>surface<br>area unit |
| --- | --- | --- | --- | --- | --- | --- | --- |
| 250nm | 1µg | 4.01476x10<sup>12</sup> | 1mg | 4.9x10<sup>10</sup>/mg | 81.9 | 0.2µm<sup>2</sup> | 409.5/µm<sup>2</sup> |
| 1µm | 1µg | 4.01476x10<sup>12</sup> | 1mg | 1.8x10<sup>9</sup>/mg | 2230.4 | 3.14µm<sup>2</sup> | 710.3/µm<sup>2</sup> |

### Force application to the TCR promotes TCR signalling and CD4^+^ T cell activation

We chose to utilise the Nr4a3 tocky^29^ mouse as an NFAT reporter for TCR signalling, since NFAT is necessary and sufficient for Nr4a3 expression. Briefly, the Nr4a3 reporter protein in Nr4a3 tocky exists in a blue fluorescent form with a half-life of approximately 4 hours, that ultimately decays to a longer lived red fluorescent form with a half-life of approximately 120 hours in non-dividing T cells^29–31^. These properties mean that expression of Nr4a3-blue reports active TCR signalling, whilst expression of Nr4a3-red reports on past TCR signals, and therefore we focused our studies largely on blue^+^ red^-^ forms of the Nr4a3 reporter protein (i.e., newly induced TCR signals from cells displaying no evidence of TCR signalling in the previous 120hrs, henceforth simply referred to as Nr4a3 expression).

We focused our initial studies on whether force application to membrane bound, anti-CD3ε functionalised MNPs could promote active TCR signalling and induce expression of Nr4a3. We found that dynamic force application at a frequency of 1Hz for 1 hour to both 250nm and 1µm anti-CD3ε MNPs induced significant upregulation of Nr4a3 in Nr4a3 Tocky reporter mice (Figure 2A-2B). Representative flow plots displaying Nr4a3-Blue expression from experiments utilising either 250nm or 1μm MNPs are shown in figure 2A. Alongside testing the effect of force application in the non-TCR transgenic Nr4a3 reporter mouse, we also investigated whether this finding could be replicated in transgenic mouse models, utilising the Tg4 transgenic line which expresses the Tg4 TCR specific for the MBP Ac1-9 peptide^32^. Importantly, we found that force application to CD4^+^ T cells isolated from both Tg4 transgenic mice, and Tg4 transgenic mice bred on a Rag2^-/-^ background (known as RTO mice) was able to induce upregulation of surface activation markers CD69 and CD25 in Tg4 (Figure 2C) or RTO mice (Figure 2D), when measured by flow cytometry 4 hours post force application. These data indicate that, as expected, force application to pan-TCR targeting MNPs can upregulate TCR signalling in non-transgenic TCR reporter mice (Nr4a3 Tocky), as well as TCR transgenic mouse models such as the Tg4 mouse and its Rag2^-/-^ counterpart (RTO).

**Figure 2.**
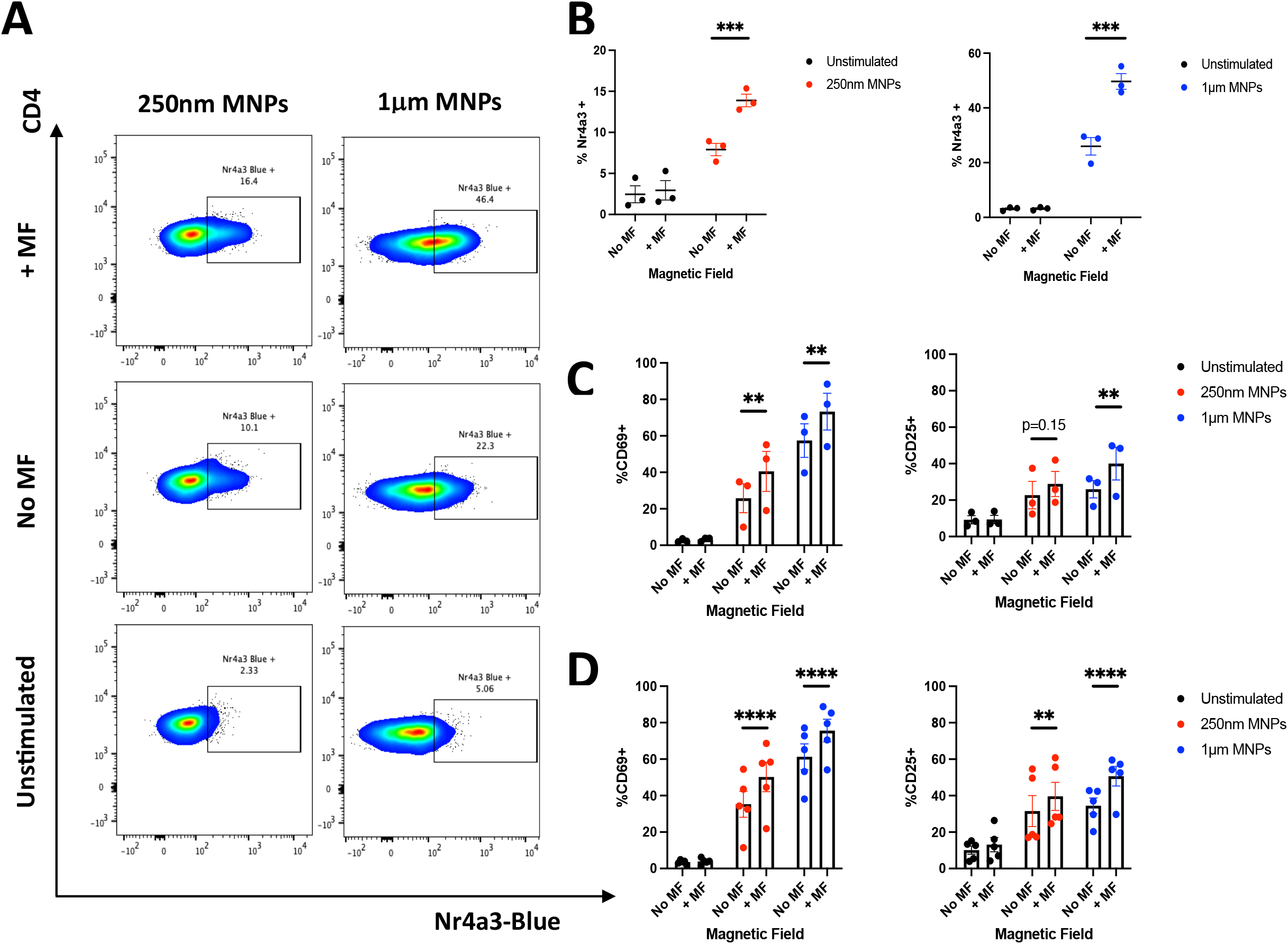
Force application for 1 hour promotes TCR signalling events in Nr4a3 Tocky reporter (A, B), Tg4 transgenic (C) and RTO mouse models (D). 250nm or 1μm MNPs functionalised with anti-CD3ε antibodies were bound to purified CD4+ T cells (Nr4a3 Tocky and Tg4 experiments), or bulk splenocytes (RTO experiments) before the application of magnetic force for 1 hour. Unstimulated controls received medium alone (i.e., no MNPs). Following incubation for 4 hours to allow for TCR signalling events to occur, cells were analysed for expression of either Nr4a3 from Nr4a3 Tocky mice (A and B), or CD69 and CD25 from Tg4 (C) or RTO (D) mice. All data shown is from a minimum of 3 independent experiments. Plots in A show representative flow cytometry plots from experiments utilising either 250nm or 1μm anti-CD3ε functionalised MNPs. Significance assessed by Two-way ANOVA with Sidak’s post tests, ** p<0.01, *** p<0.001, **** p<0.0001.

Next, we confirmed that these responses were indeed specific to signalling through the TCR by two separate methodologies. Firstly, pre-treatment of CD4^+^ T cells from Tg4 mice with the pan-SRC kinase inhibitor PP2^33^ prevented the expression of the TCR activation marker CD69 when treated with anti-CD3ε functionalised MNPs in the presence or absence of external magnetic forces (figure 3A). However, whilst encouraging that blockade of the kinases responsible for TCR signalling prevented signalling in response to CD3ε targeted MNPs, this does not rule out the possibility that force applied to these membrane bound particles was influencing TCR signalling by other means (for instance, inducing perturbations on the membrane permitting calcium entry known to be required for robust TCR signalling^34,35^). To address this, we repeated these experiments in Nr4a3 Tocky mice with off-target MNPs (MNPs functionalised with anti-MHC-I antibodies) and hypothesised that these particles should not induce evidence of TCR signalling if indeed the effect of magnetic force application was specific to targeting of the TCR-CD3 complex. Encouragingly, we found no evidence for Nr4a3, CD69 or CD25 expression when utilising anti-MHC-I targeting 250nm (figure 3B) or 1μm (figure 3C) MNPs in the presence or absence of magnetic force application. Taken together, these results indicate that force induced upregulation of TCR signalling is indeed specific to targeting of the TCR-CD3 complex.

**Figure 3.**
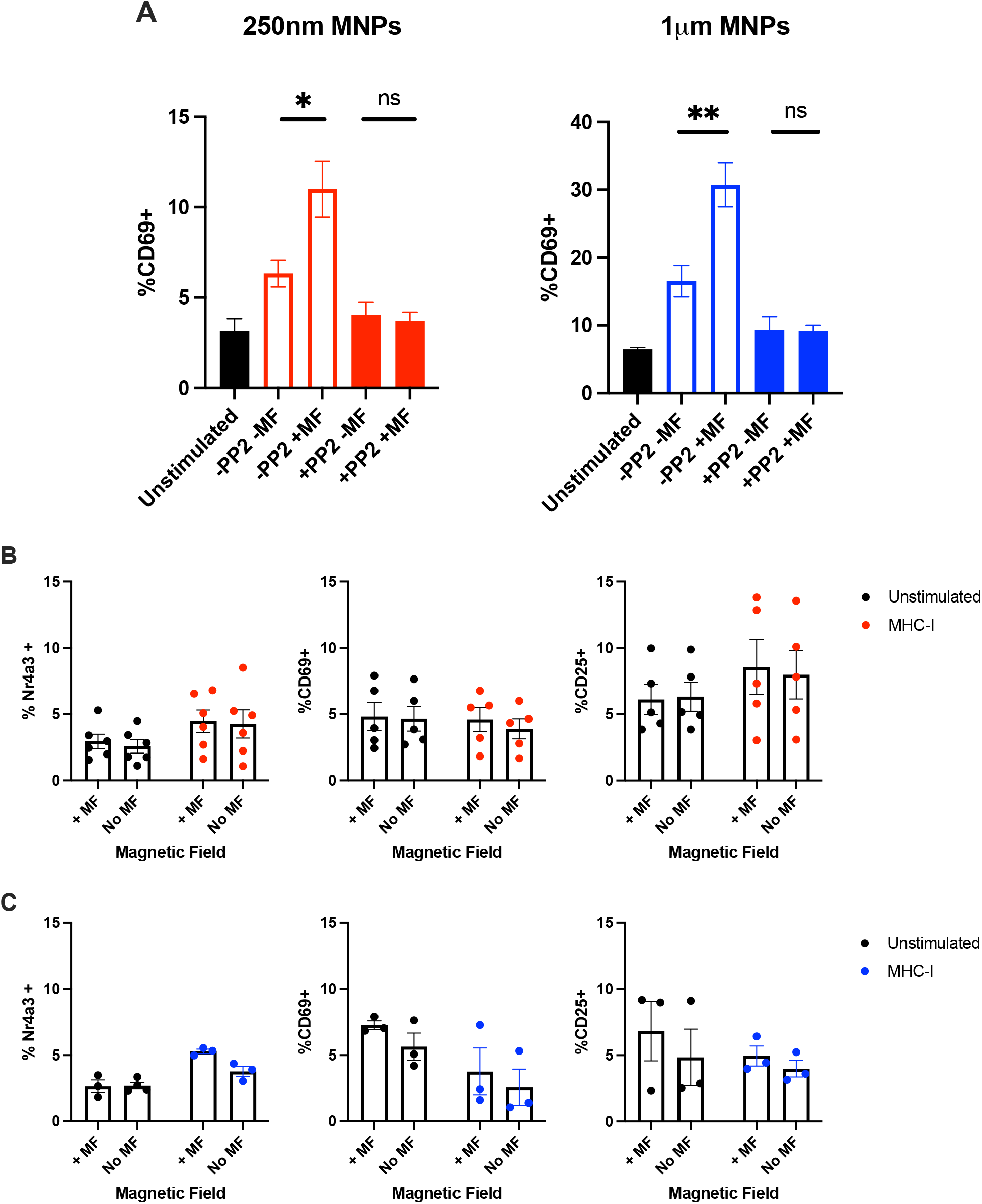
Upregulation of TCR signalling in response to force application is specific to targeting of the TCR-CD3 complex. A) Purified CD4^+^ T cells from Tg4 mice were treated with 250nm or 1µm anti-CD3ε MNPs, with or without pre-treatment with 10µM PP2, and were subjected to 1 hour of magnetic force as indicated. Pre-incubation with 10µM PP2 was sufficient to prevent upregulation of CD69 expression in response to force application. B+C) Purified CD4^+^ T cells from Nr4a3 Tocky reporter mice were treated with either 250nm (B) or 1µm (C) anti-MHC-I functionalised MNPs and subjected to 1 hour magnetic force application, before assessment of T cell activation marker expression 4 hours later. Use of anti-MHC-I targeted MNPs failed to induce expression of Nr4a3, CD69 or CD25. Data shown from a minimum of 3 independent experiments from A (250nm MNP), B and C, or from 2 independent experiments containing a total of 3 biological replicates (A, 1µm MNP). Significance assessed by one-way ANOVA where mean of each column was compared with every other column. * p<0.05, ** p<0.01.

Utilising two different sized MNPs allowed for the application of different magnitudes of magnetic force within the same magnetic field (table 1). Therefore, we were able to ask whether utilisation of the 1µm MNP was able to increase TCR signalling to a greater degree than that of the 250nm MNP. Interestingly, whilst TCR signalling in the response to the 1µm MNP was stronger when compared with the 250nm MNP (figure 2, Nr4a3, CD69 and CD25 expression, 250nm vs 1µm), this appeared not to be attributed to the strength of magnetic force application. As indicated in supplementary figure 2, when both particles were functionalised with equivalent amounts of anti-CD3ε antibody, fold-increases in Nr4a3, CD69 and CD25 expression in response to magnetic force application was highly similar for both MNPs, indicating that an increase in force application to the TCR in these settings through the use of a larger MNP does not lead to a larger increase in TCR signalling and T cell activation.

Of note is the observation that despite the magnetic force induced fold-increase in signalling being highly similar, use of the 1µm MNP induces greater activation marker expression both with and without magnetic force application when compared to the 250nm MNP, an effect potentially resulting from the differential arrangement of these antibodies on the particle surface as a result of the different particle sizes (table 2, antibody molecules/µm^2^).

### Force manipulation of TCR-CD3, but not CD28 influences TCR signalling

To this point, we have utilised MNPs functionalised with anti-CD3ε only. Therefore, we aimed to assess whether we could improve on the observed levels of TCR signalling in response to 250nm MNPs by providing co-stimulatory signalling with anti-CD28 antibodies in these settings. Here, we again applied magnetic force for 1 hour to purified CD4^+^ T cells from Nr4a3 Tocky mice, treated with particles functionalised with either anti-CD3 alone or both anti-CD3 and CD28 at a 1:1 ratio (denoted CD3:CD28). Estimated densities of antibody loading onto particle surfaces are summarised in table 2. Importantly, CD4^+^ T cells were also treated with anti-CD3 MNPs alongside soluble CD28 antibodies, in order to compare the effect of both MNP bound or soluble CD28 stimulation and hence assess whether force manipulation of the CD28 molecule could also impact T cell activation.

Unsurprisingly, we observed stronger Nr4a3 (figure 4A), CD69 (figure 4B) and CD25 (figure 4C) expression, together with T cell proliferation (figure 4D-E) under conditions where co-stimulation was provided. However, in line with other reports^36^ we identify that only force manipulation of the TCR-CD3 complex is able to induce an upregulation of T cell activation, evidenced through the highly similar responses achieved with CD3:CD28 functionalised particles compared with conditions where CD28 stimulation was provided through use of soluble antibody (figure 4).

**Figure 4.**
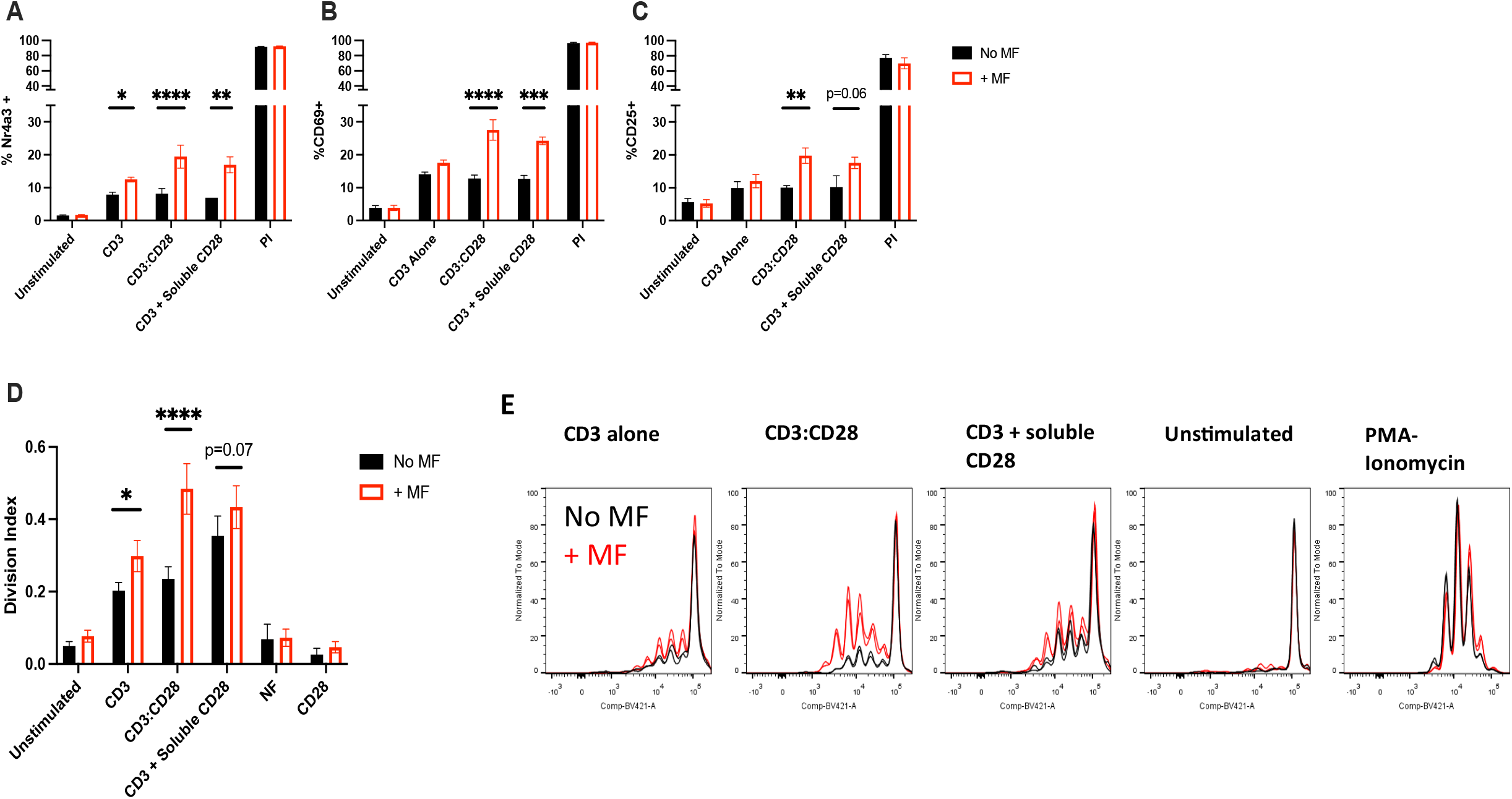
Force application to TCR-CD3, and not CD28, positively impacts T cell activation. Purified CD4^+^ T cells from Nr4a3 Tocky reporter mice were treated with 250nm MNPs functionalised as indicated, subjected to 1 hour of magnetic force, before assessment of Nr4a3 (A), CD69 (B) and CD25 (C) expression 4 hours later. Force application driven through the TCR-CD3 complex, but not CD28, positively impacts T cell signalling and proliferation (D-E) as evidenced through the comparison between MNPs coated with both anti-CD3 and anti-CD28 antibodies (CD3:CD28) and MNPs treated with anti-CD3 alone together with soluble CD28 antibody treatment. All data shown from a minimum of 3 independent experiments. E: Representative histogram traces showing cell trace violet (CTV) dilution at 3 days post force application. Statistical significance assessed via 2-Way ANOVA with Sidak’s post-tests. * p<0.05, ** p<0.01, *** p<0.001, **** p<0.0001.

### TCR downregulation occurs concurrently with CD4^+^ T cell activation

To establish a link between manipulation of the TCR and subsequent downstream signalling, we have shown that T cell activation is not achieved when either A) SRC kinases responsible for phosphorylating the TCR-CD3 complex are blocked (figure 3A), or B) T cell activation is not achieved when MNPs functionalised with antibodies against off target membrane receptors are utilised (i.e., those targeting MHC-I, figure 3B). Building on this, it has previously been shown that TCRs are subsequently downregulated and internalised as a direct consequence of TCR engagement and signalling^37–41^, which is now considered a hallmark of end-stage immune synapse formation^42^. To this end, we tracked TCR expression in response to force application, which highlighted that TCR downregulation occurs under the same conditions of force application that also mediate upregulation of T cell activation (figure 5A). Importantly, TCR downregulation was indeed prevented by pre-incubation with PP2, indicating that TCR downregulation is an active, on-going process requiring TCR signalling (figure 5B-C). Investigating TCR downregulation as a fold change induced by remote force application, we identify fold changes in TCR downregulation of 3.2 and 4.8 for 250nm and 1μm anti-CD3ε MNPs respectively (supplementary figure 3). Importantly, whilst TCR downregulation was observed in response to TCR/CD3 targeted MNPs alone when compared with no-particle controls, we find that when 250nm MNPs are functionalised with clonotypes of antibody targeting the TCR-CD3 complex that are typically considered ‘non-activating’, neither Nr4a3 expression nor TCR downregulation is observed in response to external force application (supplementary figure 4).

**Figure 5.**
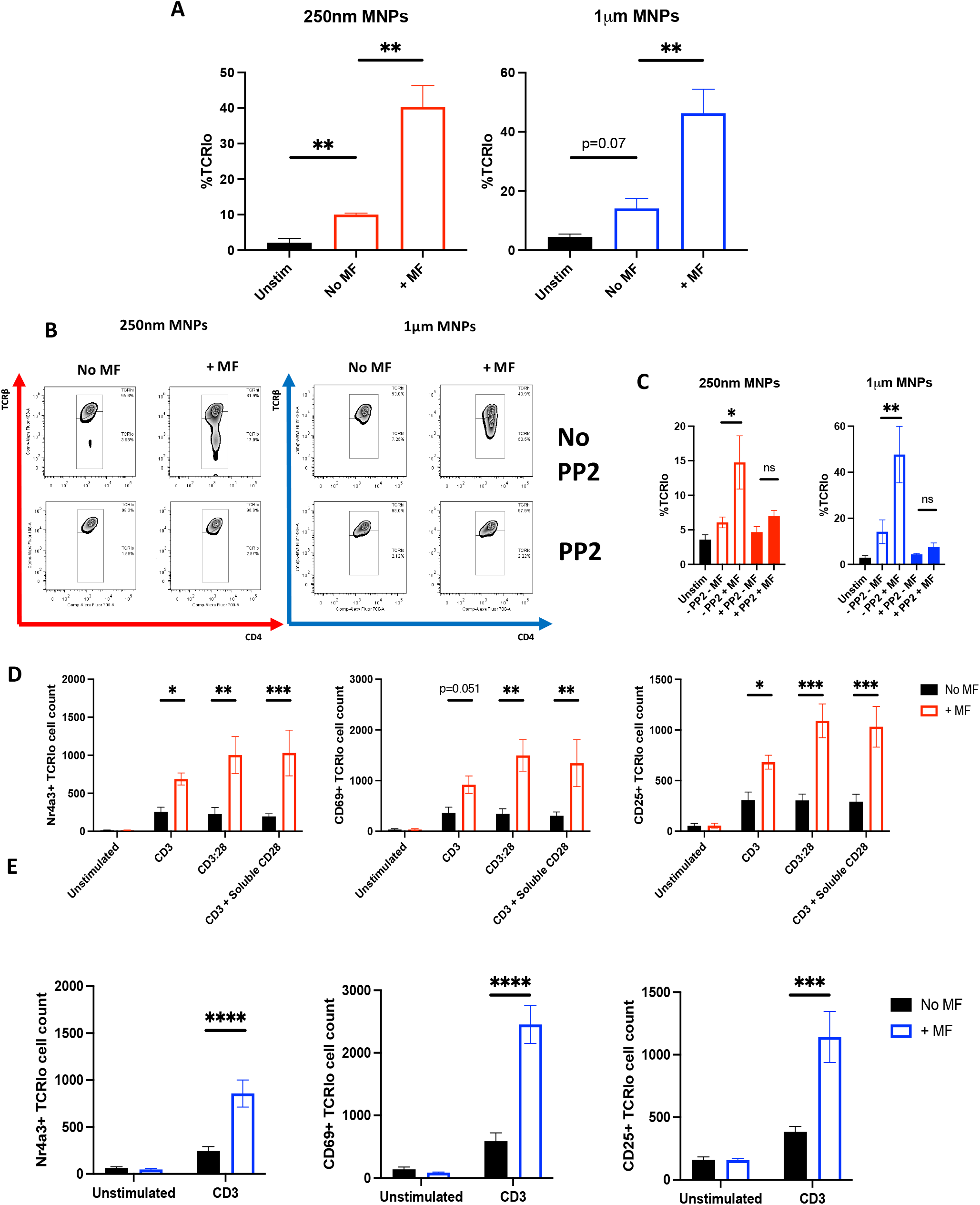
Force application to the TCR-CD3 complex induces concurrent TCR downregulation and TCR signalling. A-C: CD4^+^ T cells from Tg4 mice were treated with either 250nm or 1µm anti-CD3ε MNPs as shown, before application of magnetic force for 1 hour and assessment of TCR expression 4 hours later by flow cytometry. Cells that received neither MNP treatment or magnetic field application served as unstimulated (unstim) controls. Where indicated, T cells were pre-treated with 10µM PP2 or equivalent amount of DMSO as a control. D-E: CD4+ T cells from Nr4a3 Tocky mice were treated with 250nm or 1µm anti-CD3ε MNPs as indicated and subjected to 1 hour of magnetic force. Cells were analysed for Nr4a3, CD69, CD25 and TCR expression 4 hours later. Expression of these 3 activation markers are shown when gating on TCR^lo^ CD4^+^ populations (i.e., CD4^+^ T cells that have downregulated their TCR and become TCR^lo^ as shown in the representative gating in B). All data shown is from a minimum of 3 independent experiments. Statistical significance was assessed as follows. A: One-way ANOVA with Dunnett’s post-tests comparing the means of each group with the no MF control. C: One-way ANOVA with Tukey’s post-tests comparing the means of all groups with the mean of every other group. D, E: Two-way ANOVA with Sidak’s post-tests (D, E). * p<0.05, ** p<0.01, *** p<0.001, **** p<0.0001.

To confirm that TCR downregulation was in fact occurring in response to force application, and to rule out the observed effects being a result of steric hinderance as a result of MNP binding, we utilised a surface vs cycling TCR staining protocol (based on previously published studies for known cycling molecules such as CTLA-4^43^) to investigate the localisation of TCR molecules upon their apparent downregulation with force application. Cycling TCR staining protocols work on the notion that should the proteins be removed and internalised from the cell surface as part of known processes during TCR recycling, then the increased incubation time with staining antibody (2 hours at 37°C as opposed to 20 minutes at 4°C for surface staining) will also fluorescently tag any internalised TCR-CD3 complexes, which can then be detected by flow cytometry. This cycling staining protocol indeed reveals an increase in TCR staining intensities for both 250nm and 1µm MNPs under conditions of applied magnetic force for 1 hour (supplementary figure 5). Taken together with the observations that intracellular staining for total TCR results in staining to a highly similar intensity compared to unstimulated controls, these results identify that TCRs are in fact internalised as a direct consequence of MNP binding and magnetic force application.

Finally, since active TCR signalling appeared to be required for TCR downregulation (figures 5B-C), we sought to address whether force application to TCR-CD3 complexes was able to induce a distinct population of CD4^+^ T cells, whereby TCR signalling is induced and ultimately feeds back to control TCR downregulation, hence producing a population of TCR^lo^ CD4^+^ T cells that display evidence of TCR signalling. Figures 5D and 5E display Nr4a3, CD69 and CD25 expression patterns in TCR^lo^ CD4^+^ T cells for 250nm and 1μm MNPs respectively. Stratifying cells in this manner identified a very clear link in TCR downregulation and active TCR signalling in response to magnetic force application, whereby significant upregulation of Nr4a3, CD69 and CD25 in response to force application was observed for both 250nm and 1μm MNPs in TCR^lo^ CD4^+^ populations. Importantly, the effects of magnetic force application were not observed in cells that remain TCR^hi^ despite magnetic force application (supplementary figure 6), indicating the emergence of a TCR^lo^, activation marker positive population of CD4^+^ T cells following force application.

### Induction of a TCR^lo^ Nr4a3^+^ phenotype requires an intact actin cytoskeleton

We have shown that magnetic force application to the TCR is able to induce concurrent TCR signalling and subsequent TCR downregulation, resulting in a TCRlo Nr4a3^+^ population of CD4^+^ T cells. Importantly, TCR downregulation has been shown to occur in late stage immune synapse formation, occurring at the central supramolecular activation centre (cSMAC) where TCRs are shuttled from peripheral and distal regions of the synapse (the p/dSMAC respectively), propelled by a retrograde flow of actin polymerisation originating in the dSMAC^44,45^. Therefore, we reasoned that perturbation of the actin cytoskeleton would prevent the induction of this TCR^lo^ Nr4a3^+^ phenotype, and hence pre-treated CD4^+^ T cells with Latrunculin A (Lat A), a known actin polymerisation inhibitor which binds and sequesters actin monomers and hence prevents their polymerisation into growing actin filaments^46–48^. Here, we again observed robust TCR downregulation with both 250nm and 1μm anti-CD3ε functionalised MNPs in response to magnetic force application, which was completely inhibited by pre-treatment of CD4^+^ T cells with as little as 0.25µM Lat A (figure 6A-B). Importantly, use of Lat A allowed us to provide further evidence for a causal link between TCR downregulation and TCR signalling in response to magnetic force application. Here, we were able to investigate whether blockade of TCR downregulation with Lat A also prevented TCR signalling, or if Lat A inhibition of TCR downregulation left signalling capacity unaffected, resulting in cells that express Nr4a3 but retain their TCR expression (and hence become TCR^hi^ Nr4a3^+^). Crucially, we observed the induction of TCR^lo^ Nr4a3^+^ phenotypes in response to force application in the absence of Lat A, yet find that Lat A treatment blocked both TCR downregulation and the induction of Nr4a3 expression in response to magnetic force treatment (i.e., in the presence of Lat A, cells did not simply shift from a TCRlo Nr4a3^+^ to a TCRhi Nr4a3^+^ phenotype) (figure 6C-D). Taken together, our data on TCR downregulation proposes that remote force manipulation of the TCR-CD3 complex is sufficient to induce TCR-CD3 phosphorylation, activation of TCR signalling pathways and subsequent downregulation of TCR-CD3 complexes.

**Figure 6.**
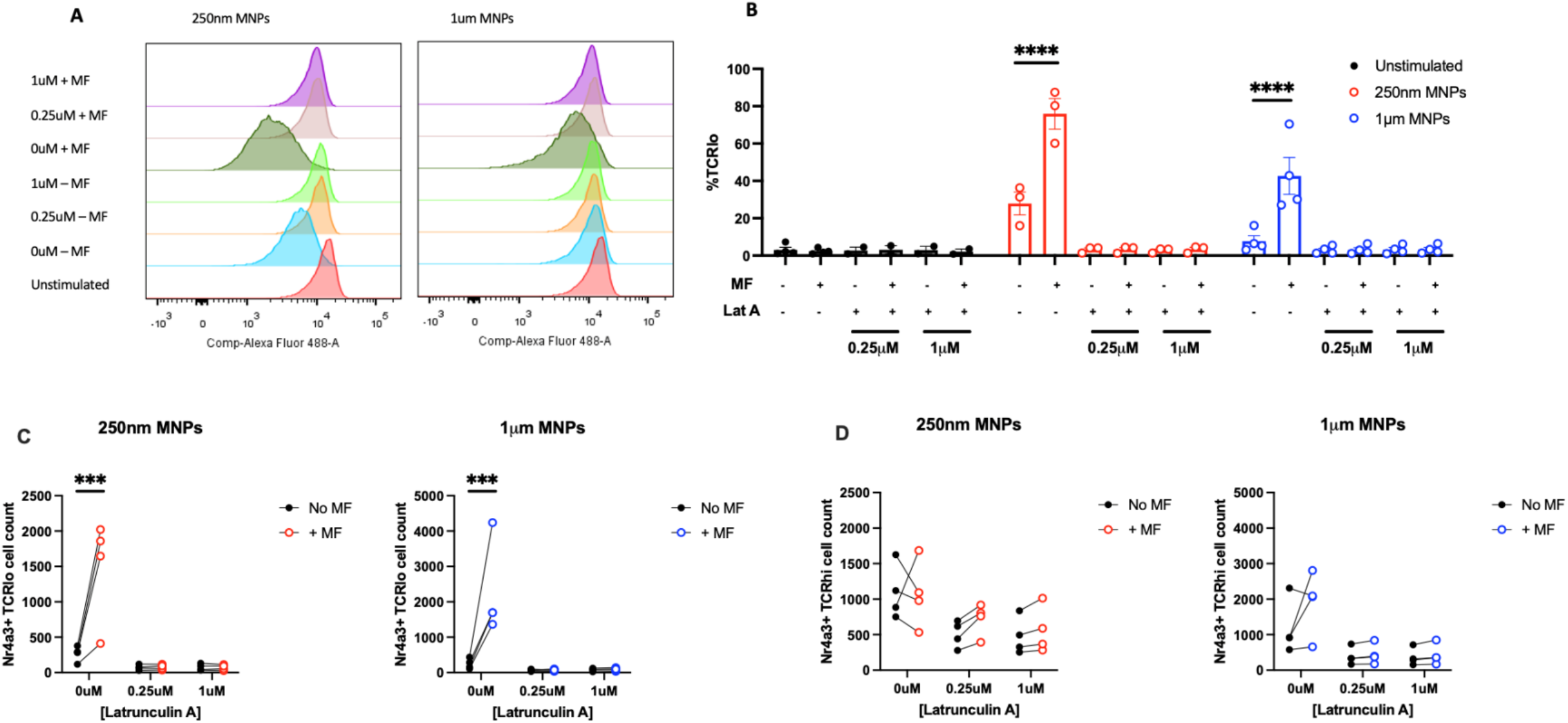
Actin cytoskeletal blockade via Latrunculin A abolishes TCR^lo^ Nr4a3^+^ populations in response to magnetic force application. CD4^+^ T cells from Nr4a3 Tocky reporter mice were treated with anti-CD3ε MNPs as indicated and subjected to 1 hour of magnetic force. Where shown, cells were pre-treated with concentrations of Latrunculin A indicated or equivalent amounts of DMSO as a control. Representative TCRβ expression profiles are shown in A. B: Percentage of cells downregulating their TCR across 3 (250nm MNPs) or 4 (1µm MNPs) independent experiments, when stimulated with either 250nm or 1µm anti-CD3ε MNPs with Latrunculin A treatment as shown. Expression patterns of Nr4a3 within either TCR^lo^ (C) or TCR^hi^ (D) populations indicate that Latrunculin A treatment prevents both TCR downregulation and Nr4a3 expression. All data shown from a minimum of 3 independent experiments. Statistical significance assessed via two-way ANOVA with Sidak’s post-tests. * p<0.05, ** p<0.01, *** p<0.001, **** p<0.0001.

### Remote force application induces TCR cluster formation within 5-15 minutes

Since direct magnetic manipulation of the TCR appeared to promote T cell signalling and activation, evidenced through the induction of TCR^lo^ Nr4a3^+^ populations in response to force application (figures 5 and 6), we sought to directly visualise the distribution of the TCR throughout the course of magnetic force application by single molecule localisation microscopy (SMLM). Representative images and TCR cluster maps from these experiments are shown in supplementary figures 7 and 8. Here, CD4^+^ T cells were treated with 1µm anti-CD3ε MNPs, and subjected to magnetic force for either 5, 15 or 30 minutes before fixation and staining for TCR visualisation. Additionally, CD4^+^ T cells with 1µm anti-CD3ε MNPs were subjected to magnetic force for 60 minutes and kept in culture for a further 2 hours (i.e., replicating the conditions of the surface vs cycling staining shown in supplementary figure 4) as TCRs were shown to be internalised by this time point. As a positive control, CD4^+^ T cells were stimulated on glass immobilised anti-CD3ε and anti-CD28 for 15 minutes to induce TCR cluster formation, which have been previously identified as the site of active TCRs^49–51^. Interestingly, force application for as little as 5-15 minutes was sufficient to increase the number of TCR clusters detected at the cell surface, after which formation of TCR clusters appeared to resolve (figure 7A-B). This data indicates that whilst force application for 1 hour is sufficient to influence T cell activation, TCR cell surface distribution is influenced on a much shorter time scale (5 – 15 minutes).

**Figure 7.**
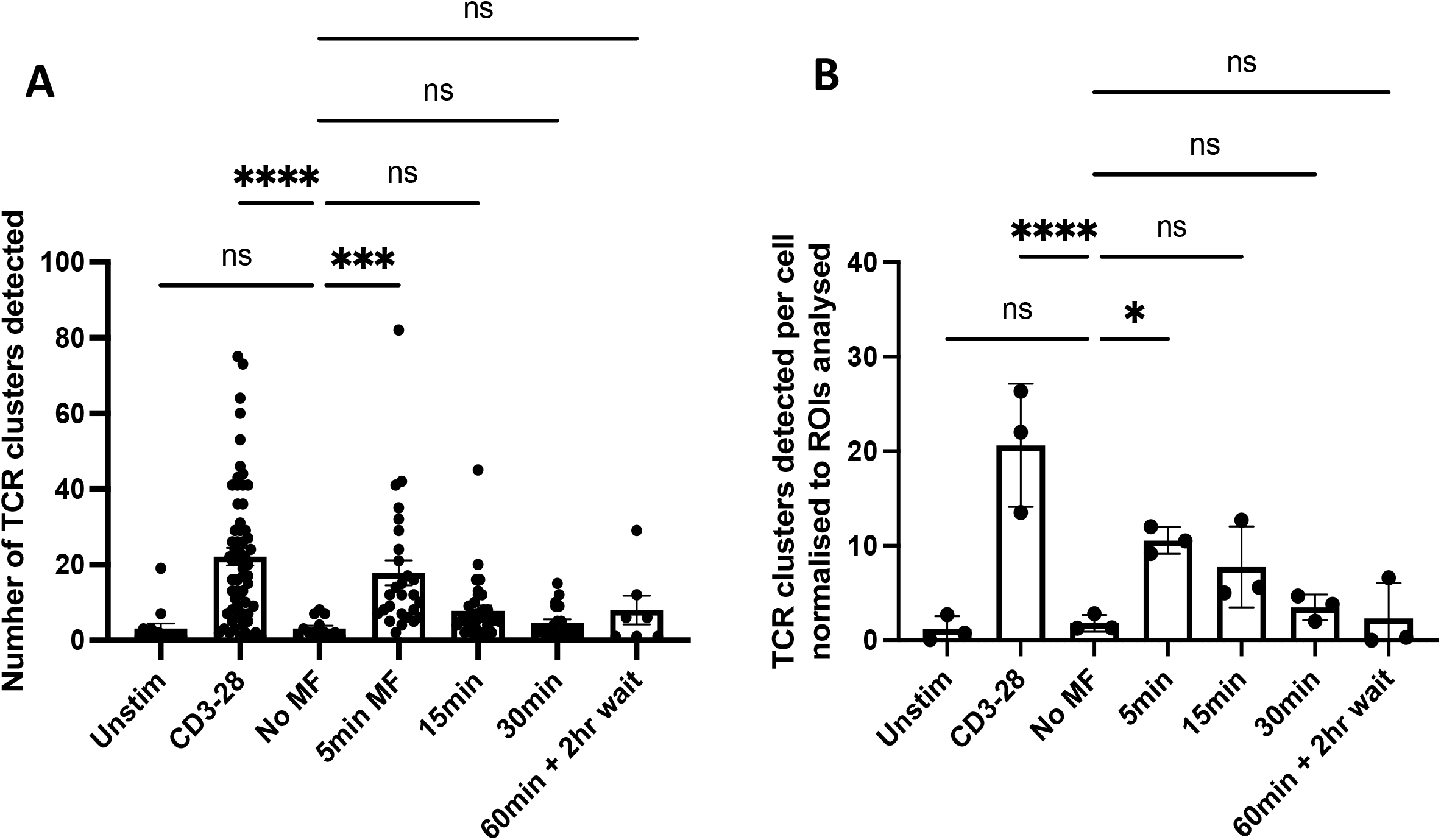
Force manipulation of the TCR-CD3 complex induces phosphorylation dependent TCR cluster formation. CD4+ T cells from Tg4 mice were treated with 1µm anti-CD3ε MNPs and subjected to magnetic force for the time frames shown, before fixation and staining for analysis of membrane TCR distribution via SMLM. Cells were stimulated on glass immobilised anti-CD3/CD28 antibodies as a positive control. Force application for 5 – 15 minutes induced the formation of membrane TCR clusters, which appear to dissociate beyond these time points. Data shown in both A and B are both from 3-4 individual independent experiments, within a minimum of 6 ROIs from individual cells analysed per condition per experiment. Statistical analysis performed via Kruskal-Wallis test with Dunn’s multiple comparison post-tests (A), or one-way ANOVA with Sidak’s multiple comparison post-test (B). A: **** p<0.0001, ***p=0.0003. B: **** p<0.0001, * p=0.0422.

### Remote force application permits the reaching of T cell activation thresholds

Since remote force application appeared to influence cell surface distribution on a much shorter time scale, we sought to understand the temporal relationship between TCR cluster formation and distal T cell signalling and activation. To investigate this, we probed whether persistent force application is sufficient to reach the signalling threshold required for T cell activation. Since T cells are known to globally integrate inputs from individual TCR-CD3 molecules both spatially and temporally^52,53^, we hypothesised that cyclical application of magnetic force for 4 rounds of 15 minute force pulses, each separated by a 15 minute break (i.e., accumulating to a total force application time frame of 60 minutes as displayed in figure 8A) might induce similar levels of Nr4a3 expression when compared to one full, uninterrupted period of 60 minutes of force application. Figure 8B shows that whilst force application for one 15-minute pulse induced a small upregulation in Nr4a3 expression (<5% increase in cells expressing Nr4a3, No MF: 14%, 15min MF: 18.8%), robust upregulation of Nr4a3 expression was only observed with either 15min x 4 or 1 uninterrupted 60 minute application of external magnetic force (%Nr4a3+: 60min: 14% no MF vs 25% +MF. 15min x 4: 15% no MF vs 32% +MF).

**Figure 8.**
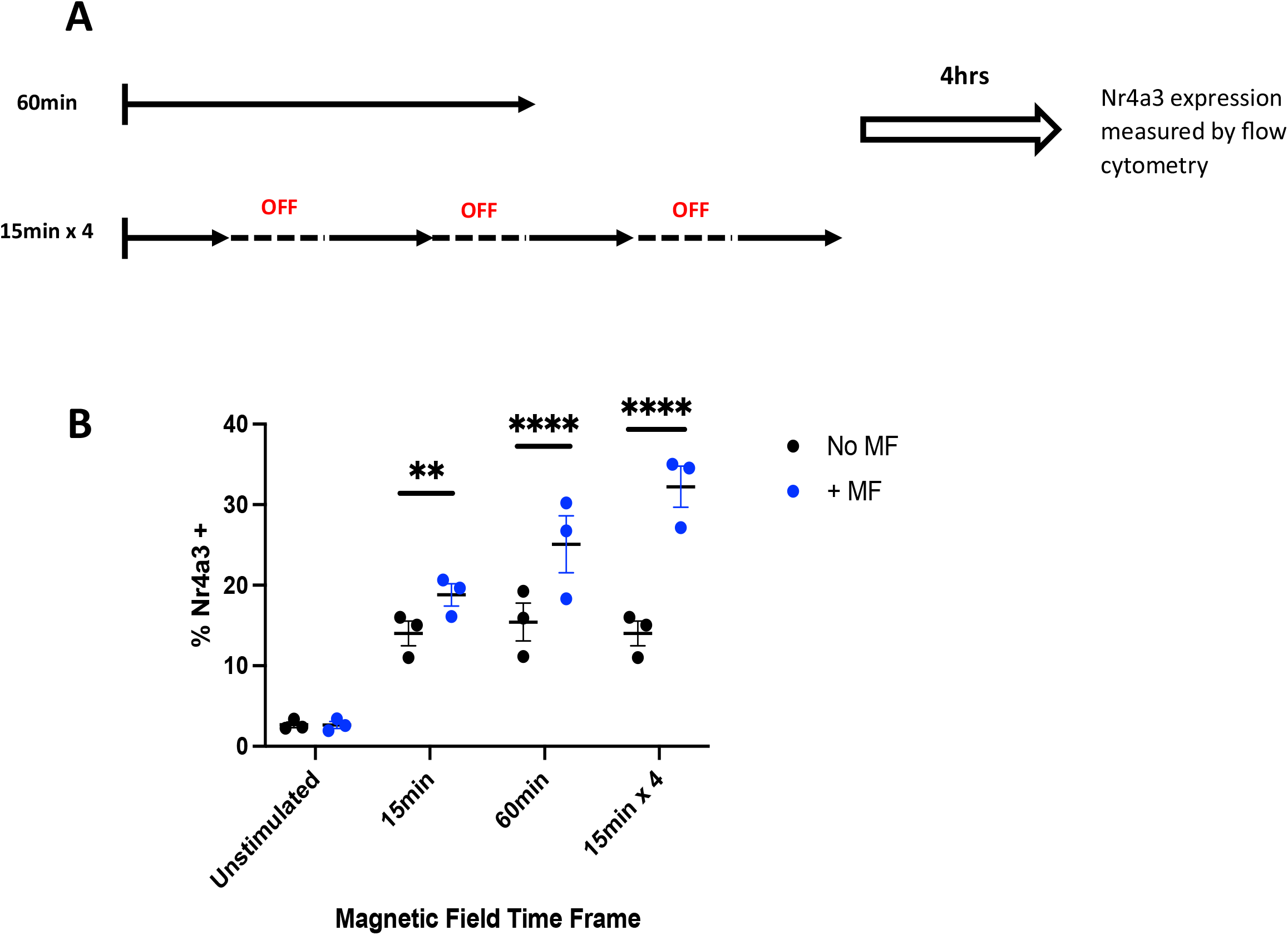
Cyclical force application induces signal accumulation along the NFAT-Nr4a3 axis, promoting Nr4a3 expression. A: Schematic representing the experimental set up. CD4^+^ T cells were treated with 1µm anti-CD3ε MNPs, before application of external magnetic force for either one full 60-minute treatment, or 4 rounds of 15 minutes of force treatment, each separated by a 15-minute period of no force application. B: Robust Nr4a3 expression is achieved with both 60 minutes of magnetic force application and 15 x 4 minutes as outlined in A. Data shown from 3 independent experiments, statistical analysis via two-way ANOVA with Sidak’s post-tests. ** p<0.01, **** p<0.0001.

Expression of Nr4a3 downstream of TCR engagement is known to be triggered through the calcineurin-calmodulin-NFAT pathway^54,55^, which converges on the dephosphorylation of NFAT permitting nuclear entry and subsequent transcription of Nr4a3. In the Nr4a3-tocky mouse, it is known that use of the calmodulin inhibitor Cyclopsorin A (henceforth called CsA) prevents Nr4a3 expression upon TCR engagement^54^. Therefore, we hypothesised that if force application is inducing an accumulation of signalling events along the NFAT pathway, blockade of the pathway by addition of 1µM CsA during each sequential break between 15 minute force pulse (figure 9A) should reveal this across the time course. As shown in figure 9B, blockade of the NFAT pathway through addition of CsA reveals a time dependent increase in Nr4a3 expression across the force application protocol, reaching significance when total time for force application reaches 60 minutes (i.e., with 1 full uninterrupted 60 minute force pulse, or when all four 15 minute pulses are completed).

**Figure 9.**
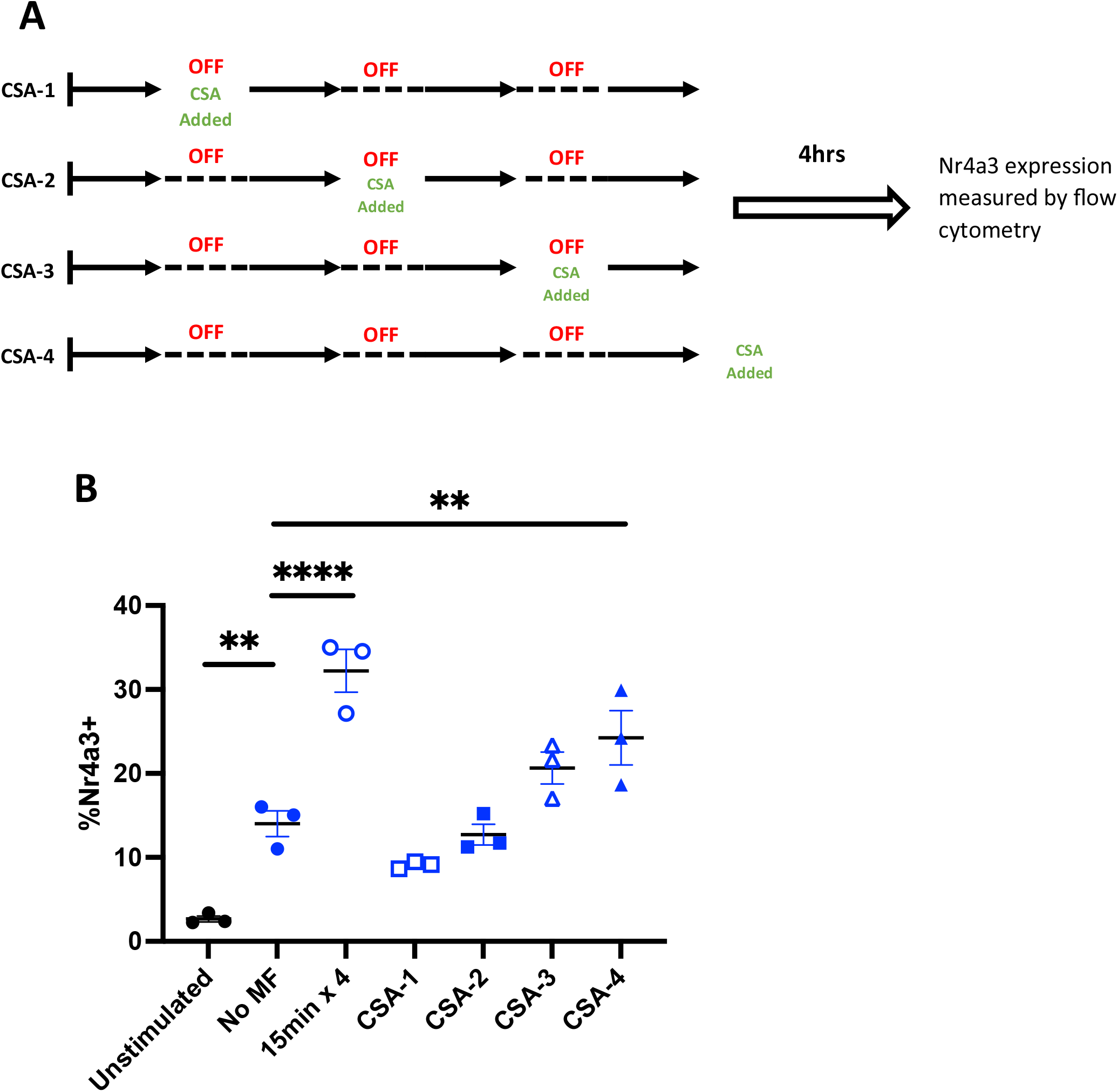
Blockade of calcineurin prevents signal accumulation along the Calcineurin-NFAT-Nr4a3 axis. CD4^+^ T cells from Nr4a3 Tocky mice were treated with 1µm anti-CD3ε MNPs and subjected to 4 rounds of 15 minute periods of magnetic force treatment, each separated by a 15 minute period of no magnetic force as per figure 7A. A: To probe signal accumulation along the NFAT signalling pathway, cells were treated with 1µM of the calcineurin inhibitor cyclosporin A after the first (CSA-1), second (CSA-2), third (CSA-3) or fourth (CSA-4) 15 minute magnetic force pulse. B: Blockade of calcineurin prevents magnetic force induced signal accumulation along the NFAT signalling pathway, preventing Nr4a3 expression. Data shown from 3 independent experiments. Statistical analysis assessed with one-way ANOVA, with Dunnett’s multiple comparison post-tests, comparing the mean of each group to the mean of the no magnetic force control. ** p<0.001, **** p<0.0001.

Interestingly, treatment with CsA failed to prevent force induced TCR downregulation (supplementary figure 9), in the manner that treatment with PP2 did (figure 5C-D). Results arising from the use of these two inhibitors of the TCR signalling cascade reveal that proximal blockade of TCR signalling (i.e., through use of PP2) is sufficient to block both Nr4a3 expression and TCR downregulation (figures 3A and 5B-C respectively). However, use the more distal signalling inhibitor CsA is sufficient to prevent force induced Nr4a3 upregulation (figure 9) but fails to prevent TCR downregulation under the same conditions (supplementary figure 9), identifying the differential signalling requirements for commitment to these processes downstream of TCR engagement.

## Discussion

We have shown that remote force application to the TCR-CD3 complex, achieved through non-invasive external magnetic fields applied to anti-CD3ε targeted superparamagnetic nanoparticles, is able to promote TCR signalling and T cell activation. We identify that force application for 1 hour is able to upregulate TCR signalling events in CD4^+^ T cells from a multitude of mouse models tested (Nr4a3 Tocky reporter mice, Tg4 transgenic mice and the Tg4 Rag2^-/-^ KO line RTO). Importantly, TCR activation observed in response to MNPs and external magnetic forces is indeed specific to TCR triggering, evidenced through inhibition of these responses by pre-incubation with the SRC-kinase inhibitor PP2 and the lack of signalling observed in response to off-target membrane-protein targeting MNPs (i.e., MHC-I targeting MNPs, figure 3).

TCR signalling experiments monitoring Nr4a3 expression identified an approximate 1.7-2-fold increase in activation marker expression in response to external force application (supplementary figure 2), regardless of the size of MNP used (250nm vs 1µm). As larger MNPs are known to elicit a greater degree of force in a magnetic field of a given field strength and gradient, we can hence rule out force magnitude as a discriminating factor in inducing TCR signalling in these experiments. We therefore aimed to assess other mechanisms that may control the TCR signalling and activation response upon force application to the TCR-CD3 complex. We readily observed TCR downregulation in the event of force application to the TCR-CD3 complex, but importantly only under those conditions that clearly mediate TCR signalling (figure 5 and supplementary figure 4, where further TCR downregulation is not achieved by magnetic force application to anti-TCR/CD3 targeting particles that do not induce TCR signalling). These findings suggest an intriguing scenario: should force application to the TCR in these settings be sufficient to induce localised TCR signalling, is subsequent remodelling of the TCR within the membrane able to promote sustained signalling required for T cell activation? It is known that during pMHC recognition and T cell activation that engaged, active TCRs are organised into TCR-microclusters (TCR-MCs)^44,49,50,56,57^, which have been identified as the site of active TCR signalling and can indeed achieve lateral propagation of the active TCR signal through the recruitment of non-engaged receptors to these regions^58^. Subsequently, it is known that during the course of T cell activation and immune synapse formation, signalling active TCR-MCs are transported from peripheral regions of the synapse (pSMAC/dSMAC) to the cSMAC^44^, propelled largely due to retrograde flow of actin polymerisation^45^, where they are ultimately internalised^59^. During this period of travel from pSMAC to cSMAC, antigen engaged TCRs interact with TCR signalling mediators, promoting signal propagation. In our experiments, we find that pre-incubation of CD4^+^ T cells with either the SRC-kinase inhibitor PP2, or perhaps more importantly the actin polymerisation inhibitor Latrunculin A, prevents the emergence of TCR^lo^ Nr4a3^+^, CD69^+^ or CD25^+^ populations in response to magnetic force application. Therefore, we propose that force application to the TCR-CD3 complex is sufficient to induce organisation of TCRs into signalling active TCR-MCs, and that signal propagation from these force induced TCR-MCs permits the reaching of T cell activation thresholds.

Finally, within this work we have interrogated the time frame required for force application to permit the reaching of such TCR activation thresholds. Surprisingly, we found that force application for a minimum of 1 hour was required to drive substantial upregulation of T cell activation markers. Interestingly, this appears to be in line with other reports. For example, studies with an optogenetically controlled CAR-T cell, whereby signalling duration from the CAR-construct can be controlled through the exposure of the CAR-T cell to light, showed that active signalling was required for a minimum of 30 minutes to drive significant upregulation of surface CD69 or NFAT mediated GFP expression^53^. Here, we reasoned that since membrane TCR clusters, upstream-hallmarks of TCR signalling, were forming on a much shorter time scale than 1 hour (5-15minutes, figure 7 and figure S7-S8), that continued downstream signalling through the NFAT-pathway is required to reach the TCR activation threshold, as measured by ‘productive’ TCR signalling in the form of Nr4a3 expression. To test this, we compared force application for 1 hour with 4 pulses of force application for 15 minutes each separated with a 15 minute period of no force application (i.e., so that the total ‘on’ time is equivalent to 60 minutes, figure 9A). We identify that signalling events induced downstream of force application display an additive pattern, reaching similar levels of Nr4a3 expression when signalling input was pulsed compared to when signalling input was received as one entire 60minute treatment of force application, suggesting the existence of memory within the TCR signalling pathway. Importantly, treatment of the CD4^+^ T cells with Cyclosporin A (a known NFAT pathway inhibitor^54^) in between each pulsatile signalling input (figure 9A) prevented this accumulation of Nr4a3 expression, proving this additive mechanism of signalling through the TCR-NFAT pathway. Considering these results in the context of T cell interactions with APCs, we speculate that the ability of T cells to remember TCR signalling inputs in the short term and activate in an additive manner may explain how T cells can activate as proposed by models such as the serial engagement model, which postulate that sequential productive interactions of a T cell with multiple APCs can lead to T cell activation^60–62^.

In conclusion, whilst other TCR triggering mechanisms are important for TCR signalling and T cell activation, we identify that force application to the TCR-CD3 complex can play a positive role in T cell activation and identify that accumulation of signalling events within the TCR-NFAT pathway can permit nuclear NFAT levels to reach the threshold required for CD4^+^ T cell activation. We identify that whilst as expected, co-stimulation by CD28 ligation increases the levels of TCR activation, only the TCR-CD3 complex displays mechanosensory behaviour and is able to interpret external mechanical cues. Considering the existing models of TCR triggering and subsequent TCR signalling, we find that force application to the TCR-CD3 complex is sufficient to induce membrane-organisation of TCR molecules into TCR-clusters, which have previously been identified to be the site of active TCRs ^44,49,50,56,57^. Given that the aggregation model of TCR signalling proposes that increased proximity of TCR molecules within one-another has the potential to increase the chance of engagement with activating kinases in the membrane, our data outlines how force mediated membrane re-organisation of TCR molecules may induce TCR triggering. How the temporal dynamics of cluster formation in the membrane overlap with the dynamics of segregation of these clusters from phosphatases such as CD45 is an important open question for future investigation. Ultimately, this study highlights the important role extracellular mechanical cues can play in controlling T cell activation and prompts the important continued future study of how extracellular mechanical cues can alter the T cell responses in both anti-tumoral responses and during the course of infection.

## Materials and Methods

### Mice

All experimental mice were maintained under specific pathogen free conditions by the Biomedical Services Unit (BMSU) at the University of Birmingham. Experiments utilising Tg4, B10.PL or RTO mice were performed in accordance with and under the UK Home Office Project Licence number P17E7ABFB, whilst experiments utilising Nr4a3-fluorescent timer (Nr4a3 Tocky) mice were performed in accordance with and under the UK Home Office Licence number P18A92E0A. Nr4a3-Tocky^29^ mice were obtained from Dr. Masahiro Ono, Imperial College London, UK.

### CD4^+^ T cell isolation and cell culture

CD4^+^ T cells were isolated from single cell splenocyte suspensions through the use of a mouse-CD4^+^ magnisort negative selection kit (Invitrogen, 8804-6821-74) as per the manufacturer’s guidelines. All cell culture was performed in complete RPMI-1640 (Sigma-Aldrich, R0883) supplemented with final concentrations of 4mM L-glutamine (Sigma-Alrich, G7513), 1000U/mL Penicillin and 1mg/ml streptomycin (Sigma-Aldrich, P4333), 20mM HEPES (Sigma-Aldrich, H0887), 50μM β-mercaptoethanol (Gibco, 31350-010) and 10% FCS (Sigma-Aldrich, F9665).

### Magnetic particle functionalisation

Functionalisation of either 250nm (micromod, #09-02-252) or 1μm MNPs (chemicell, #1402-1) was performed using carbodiimide crosslinking of antibodies to the COOH coated particle surface. Briefly, 1mg of particles were activated by addition of 20μL EDAC (Sigma Aldrich, #03449) /NHS (Sigma Aldrich, #130672) solution, to final concentrations of 5mM and 17mM respectively, under constant rotation for 1 hour. Particles were washed three times in 200μL 0.1M MES buffer (Sigma Aldrich, #M3671) by magnetic separation, before resuspension in 100μL 0.1M MES buffer and addition of 2μg total of capture antibody as per table 3. Particles and capture antibody were incubated overnight at 4 degrees, under constant rotation, before being subjected to three washes by magnetic separation in 100μL 0.1M MES buffer. Particles were then resuspended in 100μL 0.1M MES buffer, and 1μg total of target antibody was added as per table 3. It should be noted that were particles were functionalised with two antibodies (i.e., 250nm particles functionalised with both anti-CD3 and anti-CD28), 1ug total of each antibody was included in the functionalisation mix at this point. Particles with target antibody were incubated for 3 hours at room temperature under constant rotation, before free antibody was quenched via addition of 10μL 25mM glycine for 30 minutes. Functionalised particles were then washed three times by magnetic separation in 100μL 0.1% BSA, before finally being resuspended in 1mL of 0.1% BSA (producing a stock particle concentration of 1mg/mL for use in CD4^+^ T cell treatments). For non-functionalised controls, all steps described were followed but addition of antibodies was omitted.

**Table 3.** Antibodies used for functionalisation of magnetic nanoparticles.

| Particle | Target | Capture Antibody | Target antibody |
| --- | --- | --- | --- |
| 250nm anti-CD3 | CD3ε | Goat anti-Armenian Hamster IgG (AffiniPure Jackson ImmunoResearch, #127-005-099) | Anti-Mouse CD3ε (Clone 145-2C11, Invitrogen, #16-0031-82) |
| 250nm anti-CD3 | CD3$\epsilon\gamma$ | Mouse anti-Rat IgG2b (Clone MRG2b, Biolegend, #408205) | Anti-Mouse CD3$\epsilon\gamma$ (Clone 17A2, Biolegend, #100201) |
| 250nm anti-TCRβ | TCRβ | Goat anti-Armenian Hamster IgG (AffiniPure, Jackson ImmunoResearch, #127-005-099) | Anti-Mouse TCRβ (Clone H57-597, Biolegend, #1092901) |
| 250nm anti-CD3/CD28 | CD3ε and CD28 | Goat anti-Armenian Hamster IgG (AffiniPure, Jackson ImmunoResearch, #127-005-099) | Anti-Mouse CD3ε (Clone 145-2C11, Invitrogen, #16-0031-82)<br>Anti-Mouse CD28 (Clone 37.51, Invitrogen, #16-0281-82) |
| 250nm anti-MHC-I | MHC-I | Rat anti-Mouse IgG2a (Clone RMGa-62, Biolegend, #407105) | Anti-Mouse H-2K<sup>b</sup> (Clone AF6-88.5, Biolegend, #116501) |
| 1µm anti-CD3 | CD3ε | Goat anti-Armenian Hamster IgG (AffiniPure, Jackson ImmunoResearch, #127-005-099) | Anti-Mouse CD3ε (Clone 145-2C11, Invitrogen, #16-0031-82) |
| 1µm anti-MHC-I | MHC-I | Rat anti-Mouse IgG2a (Clone RMGa-62, Biolegend, #407105) | Anti-Mouse H-2K<sup>b</sup> (Clone AF6-88.5, Biolegend, #116501) |

### Calculations for number of antibodies per particle and number of antibodies per μm^2^

Assuming a 100% efficiency of antibody-particle coupling, the number of antibody molecules on the surface of either 250nm or 1μm MNPs can be estimated as follows.

- Antibody molecular weight (Hamster IgG) = 150kDa = 150,000g/mol
- Amount of antibody added (g)/150,000 = Number of moles added
- Number of moles added x avagadros constant = number of antibody molecules added
- Particle surface area = 4πr^2^
- Number of antibody molecules per surface area unit = Number of antibody molecules added/4πr^2^

### Treatment of CD4^+^ T cells with functionalised MNPs and force application with MICA

Purified CD4^+^ T cells were treated with functionalised MNPs on ice for 30 minutes in complete RPMI medium, with end cell and particle concentrations being maintained at 5×10^5^/mL and 50μg/mL respectively. Cells were then washed in 20mL PBS at 300g for 5 minutes at 4°C, before cells bound with functionalised MNPs were resuspended at 1×10^6^/mL in complete RPMI for application of remote force with MICA.

For MICA force application, MNP treated CD4^+^ T cells (or relevant unstimulated controls) were plated at 4×10^5^ cells/well in a final volume of 800μL. MICA application was applied for time frames shown in figure legends throughout the body of the text. Force magnitudes applied to 250nm or 1μm MNPs were calculated as previously described ^27,28^ and are displayed in table 1.

### Flow cytometry for CD4^+^ T cell Activation

Following treatment with functionalised MNPs and force application via MICA, CD4^+^ T cells were maintained in a 37°C 5% CO_2_ incubator for a further 4 hours to allow for surface activation marker expression and for proper translation of the Nr4a3 Tocky reporter protein. Cell suspensions were then surface stained and analysed for activation marker expression, surface TCR expression and Nr4a3 Tocky reporter protein expression on an LSR-Fortessa-X20 (BD Biosciences). The following antibodies were utilised in the study; Anti-TCRβ FITC (1:200, H57-597), anti-CD4 AF700 (1:200, RM4-5), anti-CD69 PeCy7 (1:200, H1.2F3), anti-CD25 APC (1:200, PC61), fixable viability dye ef780 (1:1000, 65-0865-14, Thermo Fischer). All antibodies were from Biolegend unless stated otherwise, and were used at dilution factors shown made in FACs buffer (PBS + 2mM EDTA (Sigma-Aldrich, E7889) + 2% FCS).

### Measurement of CD4^+^ T cell Proliferation

Purified naïve CD4+ T cells were loaded with 5μM of Cell Trace Violet (CTV) proliferation dye as per manufacturers guidelines (Thermo Fischer, C34557). These cells were stimulated as described in the text, in the presence of 20U/mL exogenous IL-2 (R&D Systems, 202-IL), with proliferation being assessed by CTV dilution on an LSR-Fortessa-X20 (BD Biosciences) 72 hours post force application. Proliferation was assessed on live CD4+ T cells, achieved by gating on fixable viability dye ef780 (Cat 65-0865-14, Thermo Fischer) negative, CD4-AF700 (clone RM4-5, Biolegend) positive events. The inbuilt FloJo proliferation tool was then used to calculate the division index of each sample.

### Surface vs Cycling vs Total TCR analysis

For analysis of surface vs cycling vs total TCR analysis, purified CD4^+^ T cells were bound with functionalised MNPs as shown and subjected to 1 hour of magnetic force. For cycling TCR analysis, cells were then immediately surface stained with 100μL anti-TCRβ (H57-597, 1:200 in complete RPMI) for 2 hours at 37°C. After this, cells were washed three times in FACs buffer, and surface stained for CD4 and fixable viability dye only, before fixation with BD cytofix for 20 minutes at 4°C. For surface TCR analysis, cells were kept in complete medium alone for 2 hours following magnetic force application and were then surface stained for TCRβ, CD4 and expression together with fixable viability dye as previously described and before fixation. Finally, for TCR analysis, cells were again kept in complete medium alone for 2 hours post magnetic force application and were then surface stained and fixed as per the surface TCR expression samples. These cells were then permeabilised with 1X permeabilisation buffer (Invitrogen, 00-8333-56) at 4°C for 15 minutes before intracellular staining with anti-TCRβ.

### Single Molecule Localisation Microscopy

CD4^+^ T cells purified from Tg4 mice were treated with 1μm anti-CD3ε MNPs as described, before being subjected to magnetic force for time frames shown in glass bottomed ibidi imaging chambers (Ibidi, 80807). For positive or negative controls, cells were stimulated on glass bound anti-CD3/CD28 (2μg/mL and 1μg/ml respectively, CD3: Invitrogen, 145-2C11, #16-0031-82, CD28: Invitrogen, 37.51, #14-0281-82) for 15 minutes or were plated in non-coated chambers respectively. Following stimulation, cells were immediately fixed by the addition of PFA to a final concentration of 4% at 4°C for 20 minutes. Cells were then washed three times in 250μL PBS and blocked for 1 hour at room temperature in 250μL 5% BSA in PBS. Cells were then washed in PBS a further three times, before staining for 1 hour at room temperature with 250μL anti-TCRβ AF647 (H57-597, Biolegend, 1:100 in PBS). Cells were then washed three times in PBS, and stored overnight in 250μL PBS and imaged the following day. On the day of imaging, all conditions were put into 200μL 0.2M mercaptoethilamine (MEA, pH 7.5) prior to image acquisition.

Images were acquired on a Nikon dSTORM SMLM microscope at 100X magnification under oil immersion. Image acquisition parameters were as follows; initial fluorophore activation was achieved through exposure to the 405nm laser at 100% power for 100 frames followed by collection of fluorophore locations for 5000 frames, with the 640nm laser at 20%. This cycle was repeated for a total of 6 times, producing an acquisition period of approximately 30,000 frames and approximately 10 minutes. Images were not collected whilst fluorophore reactivation phases with the 405nm laser were taking place.

Post acquisition, all images were analysed using the FIJI ‘ThunderSTORM’ plug-in^63^ to generate point clouds of fluorophore detections. Fluorophore detections that appeared within 20nm of one another were merged to one detection to account for multiple-blinking. Within this merged image, molecules of an intensity less than 1000 were filtered out, and regions of interest were drawn within the glass-cell contact zone to exclude noise from membrane edges. Fluorophore detection lists from the resulting images were analysed using a custom R script which measured the degree of TCR clustering by DBSCAN, using the following parameters as per previous reports^64^. Minimum detections per cluster = 10, maximum detections per cluster = 10,000, DBSCAN radius = 20, DBSCAN threshold = 3. Finally, MATLAB was used to produce TCR cluster maps to visualise membrane TCR distribution.

## Acknowledgements

JC and AEH acknowledge financial support from an EU ERC Advanced Grant DYNACEUTICS (grant no. 789119). DB is funded by an MRC career development fellowship (MR/V009052/1) and a Lister Institute of Preventative Medicine Fellowship.

## References

1. Krogsgaard, M., Juang, J. & Davis, M. M. A role for ‘self’ in T-cell activation. Semin Immunol 19, 236–244 (2007).

2. Krogsgaard, M. et al. Agonist/endogenous peptide-MHC heterodimers drive T cell activation and sensitivity. Nature 434, 238–243 (2005).

3. Štefanoví, I., Dorfman, J. R. & Germain, R. N. Self-recognition promotes the foreign antigen sensitivity of naive T lymphocytes. Nature 420, 429–434 (2002).

4. van der Merwe, P. A. Do T cell receptors do it alone? Nat Immunol 3, 1122–1123 (2002).

5. Chang, V. T. et al. Initiation of T cell signaling by CD45 segregation at ‘close contacts’. Nat Immunol 17, 574–582 (2016).

6. Davis, S. J. & van der Merwe, P. A. The kinetic-segregation model: TCR triggering and beyond. Nat Immunol 7, 803–809 (2006).

7. Chen, K. Y. et al. Trapping or slowing the diffusion of T cell receptors at close contacts initiates T cell signaling. Proc Natl Acad Sci U S A 118, (2021).

8. Mckeithan, T. W. Kinetic proofreading in T-cell receptor signal transduction. Proc Natl Acad Sci U S A 92, 5042–5046 (1995).

9. Swamy, M. ZAP70 holds the key to kinetic proofreading for TCR ligand discrimination. Nat Immunol 23, 1293–1302 (2022).

10. Tischer, D. K. & Weiner, O. D. Light-based tuning of ligand half-life supports kinetic proofreading model of T cell signaling. Elife 8, 1–25 (2019).

11. Voisinne, G. et al. Kinetic proofreading through the multi-step activation of the ZAP70 kinase underlies early T cell ligand discrimination. Nature Immunology 2022 23:9 23, 1355–1364 (2022).

12. Feng, Y. et al. Mechanosensing drives acuity of αβ T-cell recognition. Proc Natl Acad Sci U S A 114, E8204–E8213 (2017).

13. Kim, S. T. et al. The αβ T cell receptor is an anisotropic mechanosensor. Journal of Biological Chemistry 284, 31028–31037 (2009).

14. Hu, K. H. & Butte, M. J. T cell activation requires force generation. Journal of Cell Biology 213, 535–542 (2016).

15. Liu, B., Chen, W., Evavold, B. D. & Zhu, C. Accumulation of dynamic catch bonds between TCR and agonist peptide-MHC triggers T cell signaling. Cell 157, 357–368 (2014).

16. Pryshchep, S., Zarnitsyna, V. I., Hong, J., Evavold, B. D. & Zhu, C. Accumulation of Serial Forces on TCR and CD8 Frequently Applied by Agonist Antigenic Peptides Embedded in MHC Molecules Triggers Calcium in T Cells. The Journal of Immunology 193, 68–76 (2014).

17. Zhao, X. et al. Tuning T cell receptor sensitivity through catch bond engineering. Science (1979) 376, (2022).

18. Mittelheisser, V. et al. Evidence and therapeutic implications of biomechanically regulated immunosurveillance in cancer and other diseases. Nat Nanotechnol (2024) doi:10.1038/s41565-023-01535-8.

19. Markides, H., et al. Translation of remote control regenerative technologies for bone repair. NPJ Regen Med 3, (2018).

20. Rotherham, M. & Haj, A. J. El. Remote Activation of the Wnt/β-Catenin Signalling Pathway Using Functionalised Magnetic Particles. (2015) doi:10.1371/journal.pone.0121761.

21. Henstock, J. R., Rotherham, M., Rashidi, H., Shakesheff, K. M. & El Haj, A. J. Remotely Activated Mechanotransduction via Magnetic Nanoparticles Promotes Mineralization Synergistically With Bone Morphogenetic Protein 2: Applications for Injectable Cell Therapy. Stem Cells Transl Med 3, 1363–1374 (2014).

22. Gonçalves, A. I. et al. Triggering the activation of Activin A type II receptor in human adipose stem cells towards tenogenic commitment using mechanomagnetic stimulation. Nanomedicine 14, 1149–1159 (2018).

23. Perica, K. et al. Magnetic field-induced t cell receptor clustering by nanoparticles enhances t cell activation and stimulates antitumor activity. ACS Nano 8, 2252–2260 (2014).

24. Hughes, S., Dobson, J. & El Haj, A. J. Magnetic targeting of mechanosensors in bone cells for tissue engineering applications. J Biomech 40, 96–104 (2007).

25. Henstock, J. R., Rotherham, M. & El Haj, A. J. Magnetic ion channel activation of TREK1 in human mesenchymal stem cells using nanoparticles promotes osteogenesis in surrounding cells. J Tissue Eng 9, (2018).

26. Hughes, S., McBain, S., Dobson, J. & El Haj, A. J. Selective activation of mechanosensitive ion channels using magnetic particles. J R Soc Interface 5, 855–863 (2008).

27. Pankhurst, Q., Connolloy, J., Jones, S. & Dobson, J. Applications of magnetic nanoparticles in biomedicine. J Phys D Appl Phys 36, R167–R181 (2003).

28. Del Sol-Fernández, S. et al. Magnetogenetics: remote activation of cellular functions triggered by magnetic switches. Nanoscale 14, 2091–2118 (2022).

29. Bending, D. et al. A timer for analyzing temporally dynamic changes in transcription during differentiation in vivo. Journal of Cell Biology 217, 2931–2950 (2018).

30. Bending, D. et al. A temporally dynamic Foxp3 autoregulatory transcriptional circuit controls the effector Treg programme. EMBO J 37, (2018).

31. Elliot, T. A. E. et al. Antigen and checkpoint receptor engagement recalibrates T cell receptor signal strength. Immunity 54, 2481–2496 (2021).

32. Liu, G. Y. et al. Low avidity recognition of self-antigen by T cells permits escape from central tolerance. Immunity 3, 407–415 (1995).

33. Hanke, J. H. et al. Discovery of a novel, potent, and Src family-selective tyrosine kinase inhibitor: Study of Lck- and FynT-dependent T cell activation. Journal of Biological Chemistry 271, 695–701 (1996).

34. Christo, S. N. et al. Scrutinizing calcium flux oscillations in T lymphocytes to deduce the strength of stimulus. (2015) doi:10.1038/srep07760.

35. Le Borgne, M. et al. Real-Time Analysis of Calcium Signals during the Early Phase of T Cell Activation Using a Genetically Encoded Calcium Biosensor. The Journal of Immunology 196, 1471–1479 (2016).

36. Judokusumo, E., Tabdanov, E., Kumari, S., Dustin, M. L. & Kam, L. C. Mechanosensing in T lymphocyte activation. Biophys J 102, L5–L7 (2012).

37. San José, E., Borroto, A., Niedergang, F., Alcover, A. & Alarcón, B. Triggering the TCR complex causes the downregulation of nonengaged receptors by a signal transduction-dependent mechanism. Immunity 12, 161–170 (2000).

38. Chakraborty, A. K. Lighting up TCR takes advantage of serial triggering. Nat Immunol 3, 895–896 (2002).

39. Gallegos, A. M. et al. Control of T cell antigen reactivity via programmed TCR downregulation. Nat Immunol 17, 379–386 (2016).

40. Valitutti, S., Miller, S., Cella, M., Padovan, E. & Lanzavecchia, A. Serial triggering of many T-cell receptors by a few peptide-MHC complexes. Nature 375, 148–151 (1995).

41. Lanzavecchia, A., Iezzi, G. & Viola, A. From TCR engagement to T cell activation: A kinetic view of T cell behavior. Cell 96, 1–4 (1999).

42. Alarcón, B., Mestre, D. & Martinez-Martin, N. The immunological synapse: A cause or consequence of T-cell receptor triggering? Immunology 133, 420–425 (2011).

43. Ovcinnikovs, V., et al. CTLA-4-mediated transendocytosis of costimulatory molecules primarily targets migratory dendritic cells. Sci Immunol 4, (2019).

44. Varma, R., Campi, G., Yokosuka, T., Saito, T. & Dustin, M. L. T Cell Receptor-Proximal Signals Are Sustained in Peripheral Microclusters and Terminated in the Central Supramolecular Activation Cluster. Immunity 25, 117–127 (2006).

45. Babich, A. et al. F-actin polymerization and retrograde flow drive sustained PLCγ1 signaling during T cell activation. Journal of Cell Biology 197, 775–787 (2012).

46. Fujiwara, I., Zweifel, M. E., Courtemanche, N. & Pollard, T. D. Latrunculin A Accelerates Actin Filament Depolymerization in Addition to Sequestering Actin Monomers. Curr Biol 28, 3183–3192.e2 (2018).

47. Coué, M., Brenner, S. L., Spector, I. & Korn, E. D. Inhibition of actin polymerization by latrunculin A. FEBS Lett 213, 316–318 (1987).

48. Ayscough, K. R. et al. High Rates of Actin Filament Turnover in Budding Yeast and Roles for Actin in Establishment and Maintenance of Cell Polarity Revealed Using the Actin Inhibitor Latrunculin-A. Journal of Cell Biology 137, 399–416 (1997).

49. Balagopalan, L., Raychaudhuri, K. & Samelson, L. E. Microclusters as T Cell Signaling Hubs: Structure, Kinetics, and Regulation. Front Cell Dev Biol 8, 608530 (2021).

50. Hashimoto-Tane, A. & Saito, T. Dynamic regulation of TCR-microclusters and the microsynapse for T cell activation. Front Immunol 7, 1–8 (2016).

51. Hashimoto-Tane, A. et al. Dynein-Driven Transport of T Cell Receptor Microclusters Regulates Immune Synapse Formation and T Cell Activation. Immunity 34, 919–931 (2011).

52. O’Donoghue, G. P. et al. T cells selectively filter oscillatory signals on the minutes timescale. Proc Natl Acad Sci U S A 118, (2021).

53. Harris, M. J., Fuyal, M. & James, J. R. Quantifying persistence in the T-cell signaling network using an optically controllable antigen receptor. Mol Syst Biol 17, 1–20 (2021).

54. Jennings, E. et al. Nr4a1 and Nr4a3 Reporter Mice Are Differentially Sensitive to T Cell Receptor Signal Strength and Duration. Cell Rep 33, 1–10 (2020).

55. Bending, D. & Zikherman, J. Nr4a nuclear receptors: markers and modulators of antigen receptor signaling. Curr Opin Immunol 81, 102285 (2023).

56. Lillemeier, B. F. et al. TCR and Lat are expressed on separate protein islands on T cell membranes and concatenate during activation. Nat Immunol 11, 90–96 (2010).

57. Sherman, E. et al. Functional nanoscale organization of signaling molecules downstream of the T cell antigen receptor. Immunity 35, 705–720 (2011).

58. Nieves, D. J. et al. The T cell receptor displays lateral signal propagation involving non-engaged receptors. Nanoscale 14, 3513–3526 (2022).

59. Martinez-Martin, N. et al. T Cell Receptor Internalization from the Immunological Synapse Is Mediated by TC21 and RhoG GTPase-Dependent Phagocytosis. Immunity 35, 208–222 (2011).

60. Egan, J. R., Abu-Shah, E., Dushek, O., Elliott, T. & MacArthur, B. D. Fluctuations in T cell receptor and pMHC interactions regulate T cell activation. J R Soc Interface 19, (2022).

61. Friedl, P. & Gunzer, M. Interaction of T cells with APCs: the serial encounter model. Trends Immunol 22, 187–191 (2001).

62. Hellmeier, J. et al. DNA origami demonstrate the unique stimulatory power of single pMHCs as T cell antigens. Proc Natl Acad Sci U S A 118, e2016857118 (2021).

63. Ovesný, M., Křížek, P., Borkovec, J., Švindrych, Z. & Hagen, G. M. ThunderSTORM: A comprehensive ImageJ plug-in for PALM and STORM data analysis and super-resolution imaging. Bioinformatics 30, 2389–2390 (2014).

64. Pageon, S. V. et al. Functional role of T-cell receptor nanoclusters in signal initiation and antigen discrimination. Proc Natl Acad Sci U S A 113, E5454–E5463 (2016).

