## Supplementary Figures for "Remote force modulation of the T-cell receptor reveals an NFAT-threshold for CD4^+^ T cell activation"

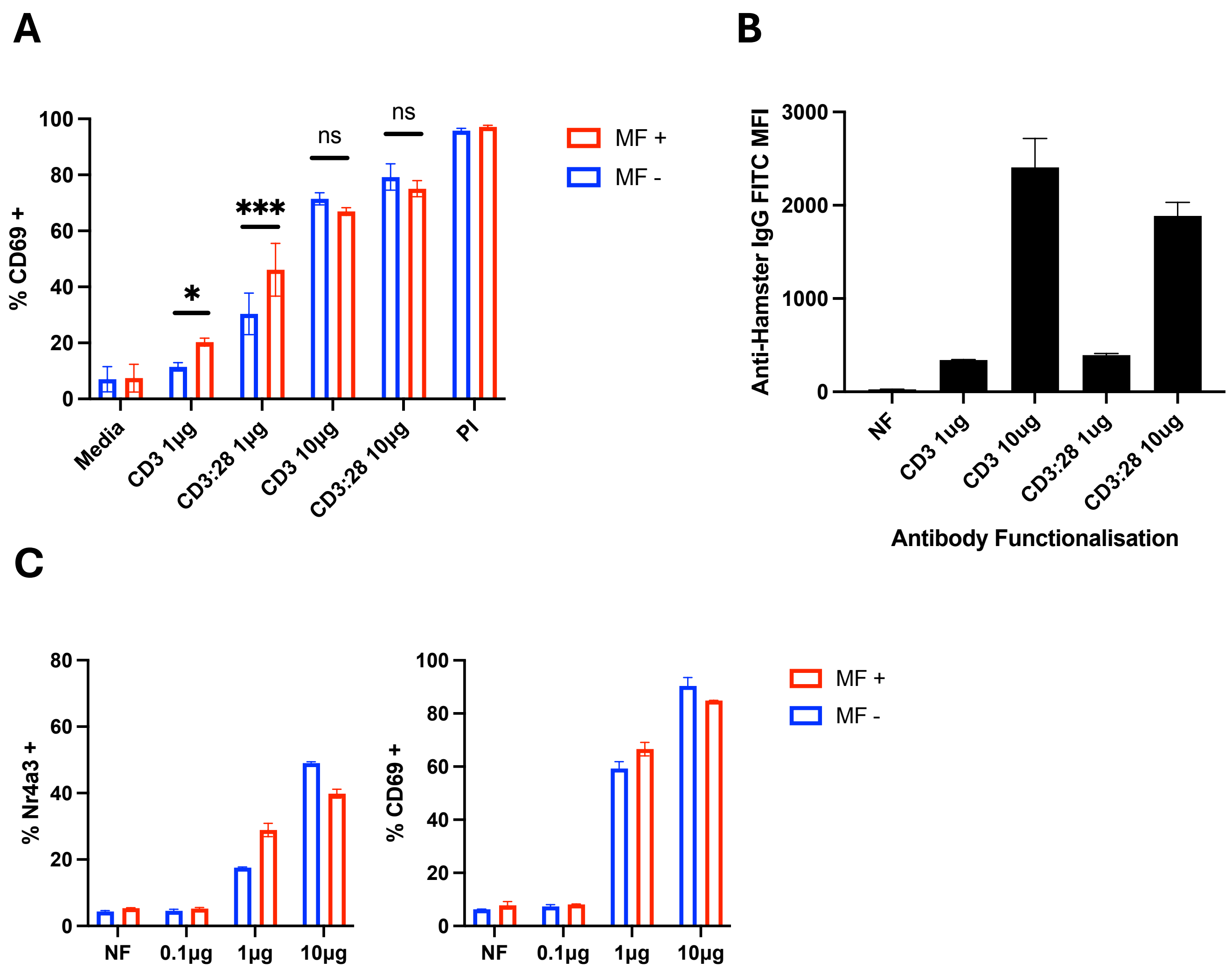


**Supplementary figure 1.** Quality control staining and functional CD4+ T cell activation experiments to optimise successful conjugation of anti-CD3 antibodies (145-2C11 clonotype) to the surface of 250nm MNPs. Particles were functionalised with the total amount of anti-CD3 or anti-CD3 and anti-CD28 antibodies as shown, as described in materials and methods. Where particles were functionalised with both anti-CD3 and anti-CD28, equal amounts of amount were included in the functionalisation mix (either 1µg of each or 10µg of each antibody). These particles were then used to activate Tg4 CD4+ T cells (A) or Nr4a3 reporter $CD4^{+}$ T cells (C), where cells were bound to particles, subject to magnetic force for 1 hour, and stained for CD69 expression 4 hours later. Functionalisation with 1µg of antibody allowed for an effect of magnetic force to be seen on T cell activation in $CD4^{+}$ T cells from Tg4 mice (CD69 expression in A) and from Nr4a3 reporter mice (Nr4a3 expression in C), whilst 10-fold increase in surface antibody increased levels of signalling observed and prevented the need for magnetic force application for T cell activation. In parallel with activation experiments, these particles were also stained with FITC conjugated anti-Armenian hamster IgG antibodies, to identify the presence of particle bound anti-CD3 antibodies. PI: PMA-Ionomycin. (B). NF: Non-functionalised. Data shown in A and B is from a minimum of 2 independent experimental repeats. Data analysis performed via two-way ANOVA with Sidak's post-tests to compare the effect of magnetic force application to each particle group. * $p=0.0101$, *** $p<0.0001$. Data shown in C is from one independent experiment, representative of 2 independent repeats.

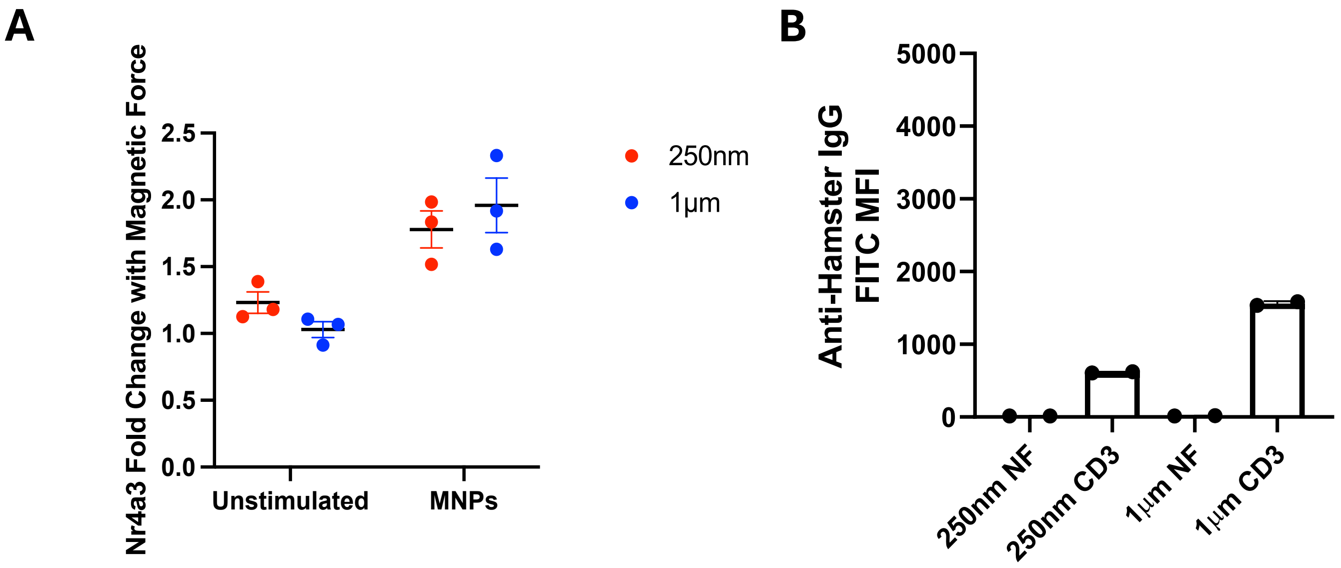


**Supplementary figure 2.** A: Fold changes in Nr4a3 expression in response to 250nm (red) or 1µm (blue) anti-CD3ε MNPs with magnetic force application. Fold changes were calculated for experiments shown in figure 2. Both 250nm and 1µm MNPs yield approximately 2-fold increase in Nr4a3 expression upon application of external magnetic force. B: MNP quality control staining for surface CD3, to assess successful conjugation of anti-CD3 to the particle surface used in experiments shown in A.

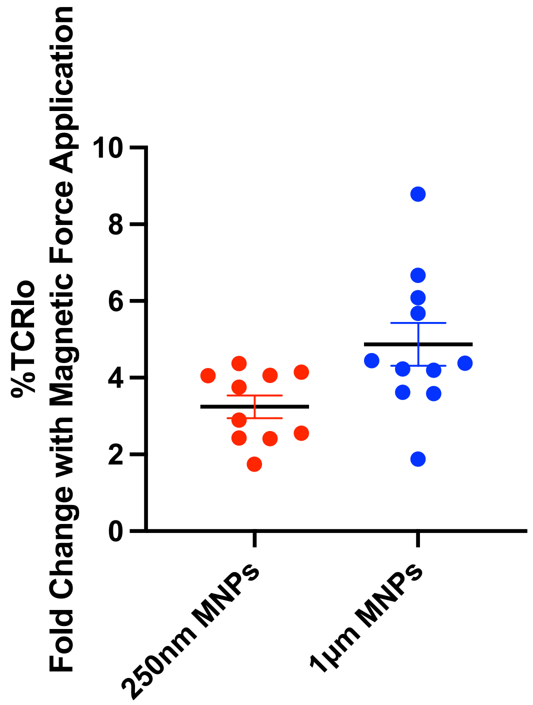


**Supplementary figure 3.** Fold change in percentage of TCRlo CD4+ T cells in response to magnetic force application to either 250nm (red) or 1µm (blue) anti-CD3ε MNPs. Fold changes were calculated for all experiments shown that analyse TCR expression. A fold change of approximately of 3.2 and 4.8 was observed for 250nm and 1µm MNPs respectively.

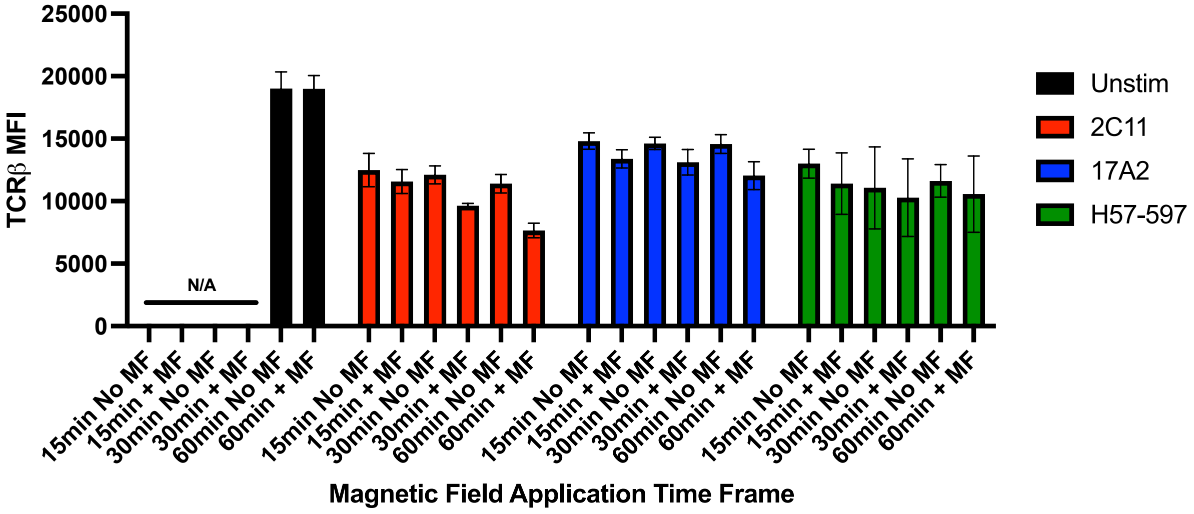


**Supplementary figure 4.** Force application to 250nm MNPs functionalised with anti-CD3 145-2C11 only induces force dependent TCR downregulation. 250nm MNPs were functionalised with either anti-CD3 145-2C11, anti-CD3 17A2 or anti-TCRβ H57-597, before force application for time frames shown and assessment of surface TCR expression 4 hours later. Unstimulated controls (no MNPs) were only treated with magnetic field for 60 minutes, or received no magnetic field treatment at all. Whilst all particle treated conditions reveal a degree of TCR downregulation compared with unstimulated controls, force application to anti-CD3 145-2C11 only induced further TCR downregulation.

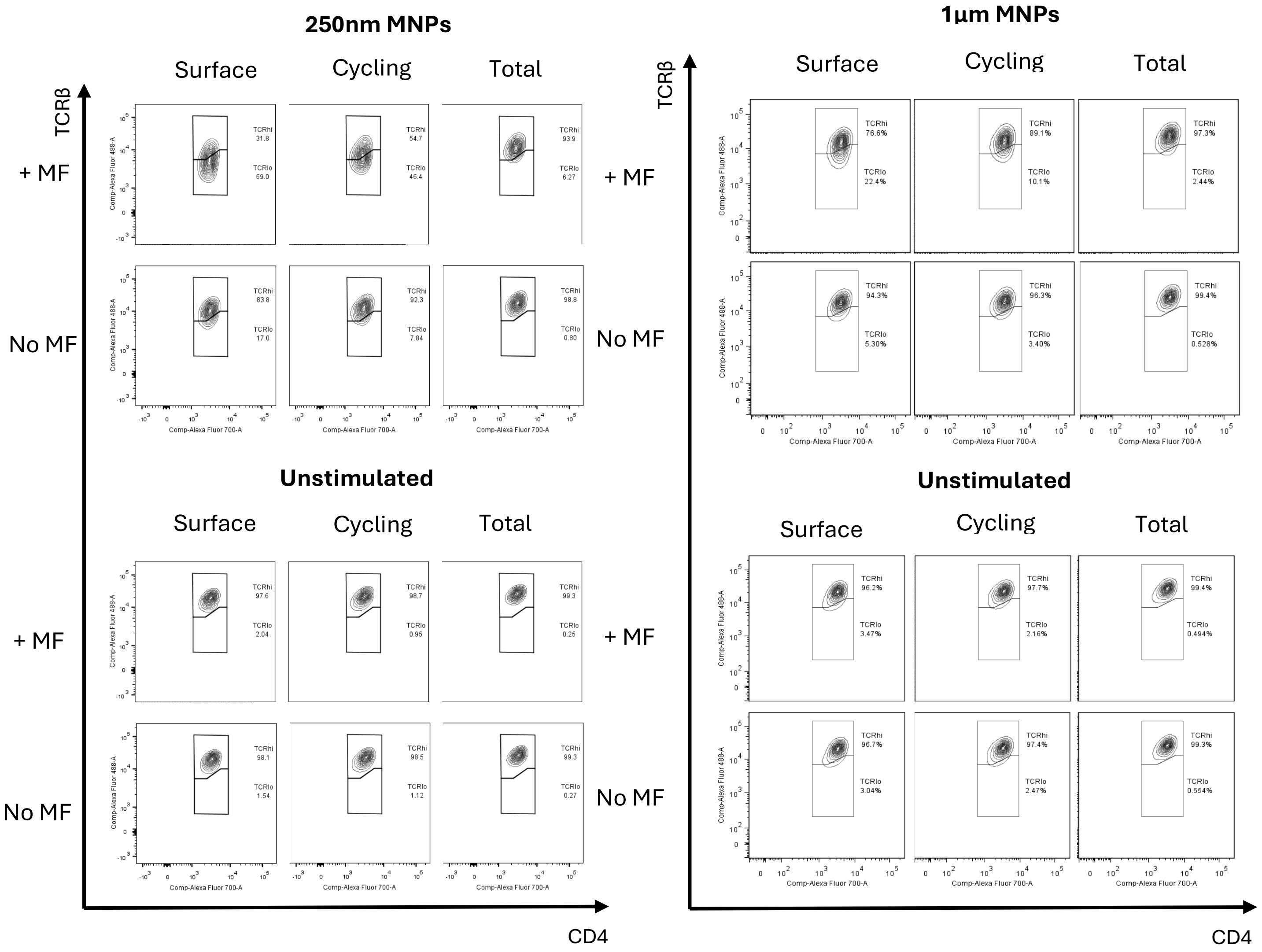


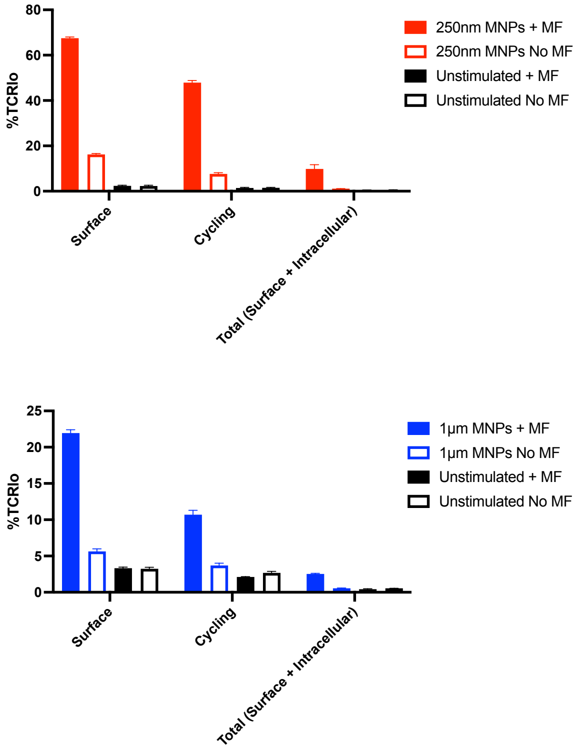


**Supplementary figure 5.** Surface vs cycling vs total TCR expression reveals that force application induces TCR internalisation. $CD4^+$ T cells treated with 250nm or 1µm anti-CD3 MNPs were subjected to 1 hour of magnetic force before staining for surface, cycling or total TCR content. Surface staining protocols alone identify a loss of surface TCR staining through application of magnetic force. Cycling staining protocols begin to show a recovery of TCR staining, which is fully recovered when staining both surface and intracellular for TCR, allowing for visualisation of TCR internalisation. All data shown from a minimum of 2 independent experimental repeats and a minimum of 3 biological repeats.

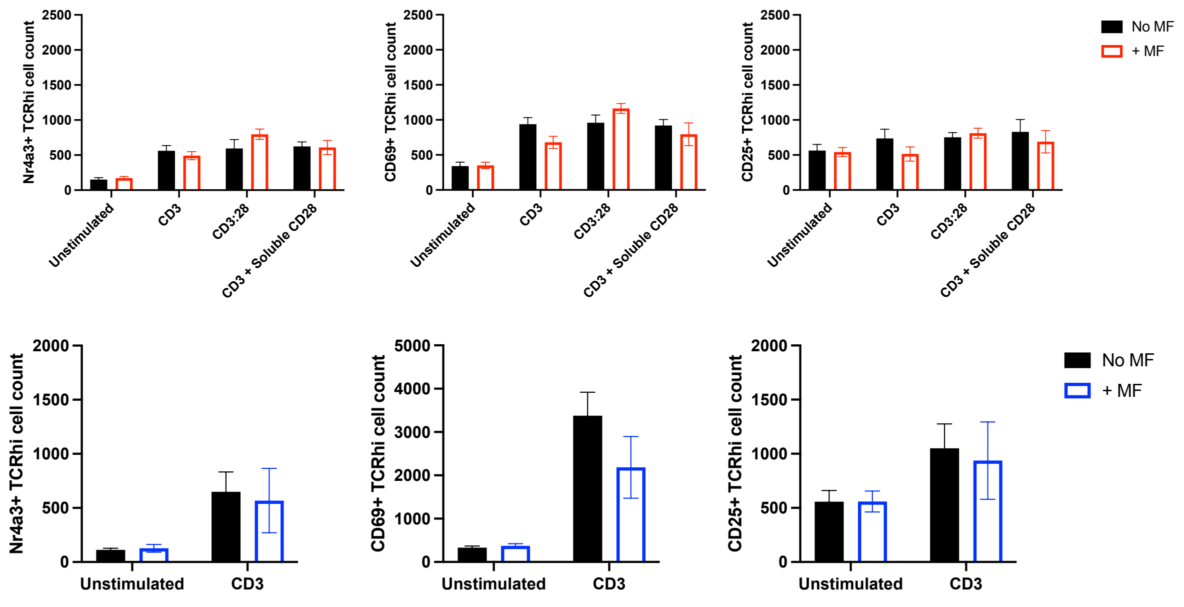


**Supplementary figure 6.** Force application to CD4⁺ T cells treated with 250nm (red) or 1µm (blue) anti-CD3 MNPs does not affect expression of T cell activation markers on cells that remain TCR$^{hi}$ when exposed to 1 hour of magnetic force. As indicated in figure 4, force application specifically induces TCR activation marker expression in cells that become TCR$^{lo}$ in response to magnetic force application. All data shown from a minimum of 3 independent experiments.

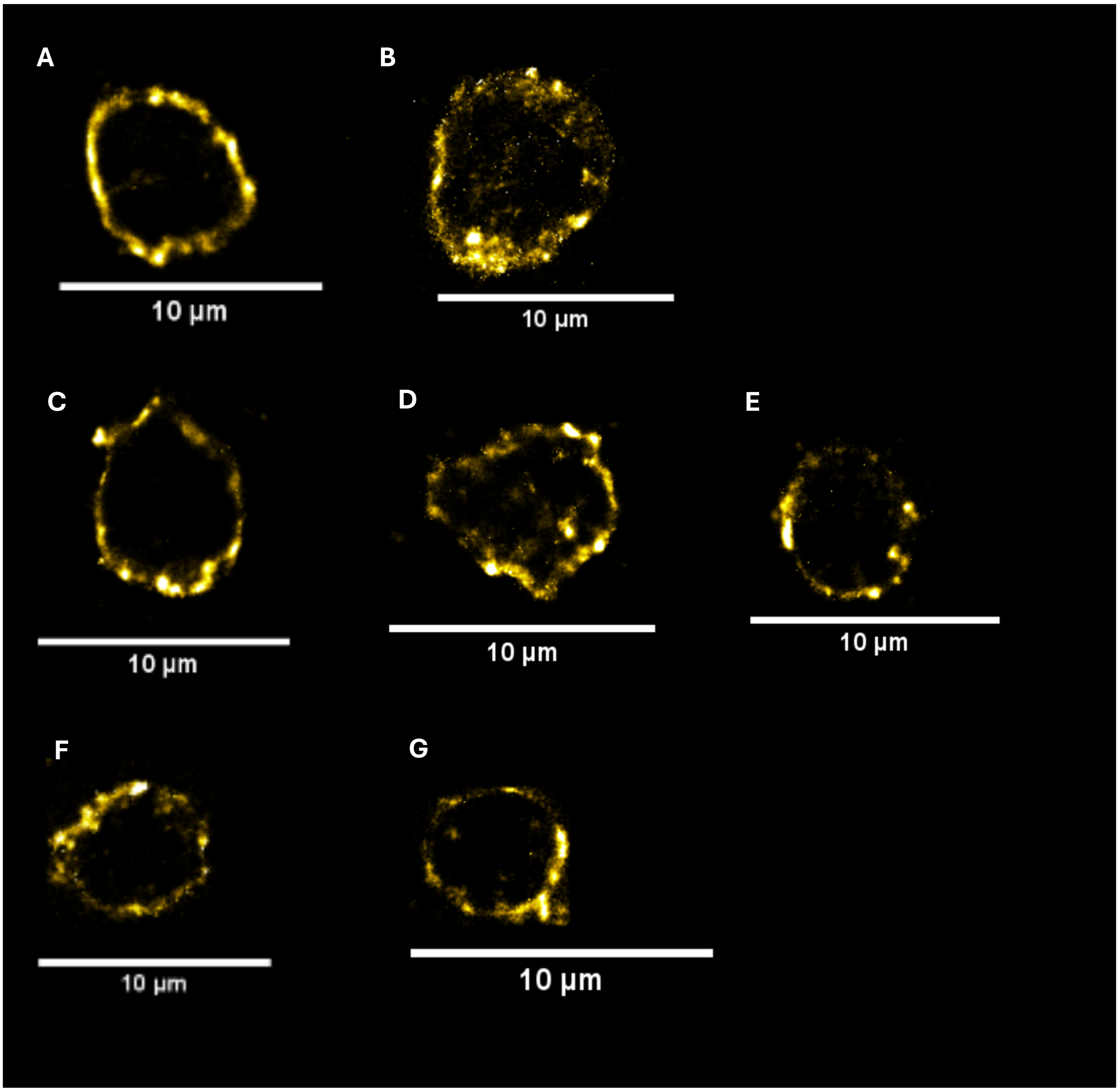


**Supplementary figure 7.** CD4$^+$ T cells from the Tg4 mouse were stimulated with 1µm anti-CD3ε MNPs as described, before being subjected to MF for 5min (D), 15min (E), or 30min (F), or 60min followed by a 2hr (G) wait as described. Cells were also either left unstimulated (A), stimulated on glass immobilised anti-CD3/CD28 (B), or were treated with 1µm MNPs but were not subjected to MF (C) as controls. Images show TCR staining from SMLM STORM imaging. Images representative of at least 2 independent experiments per condition.

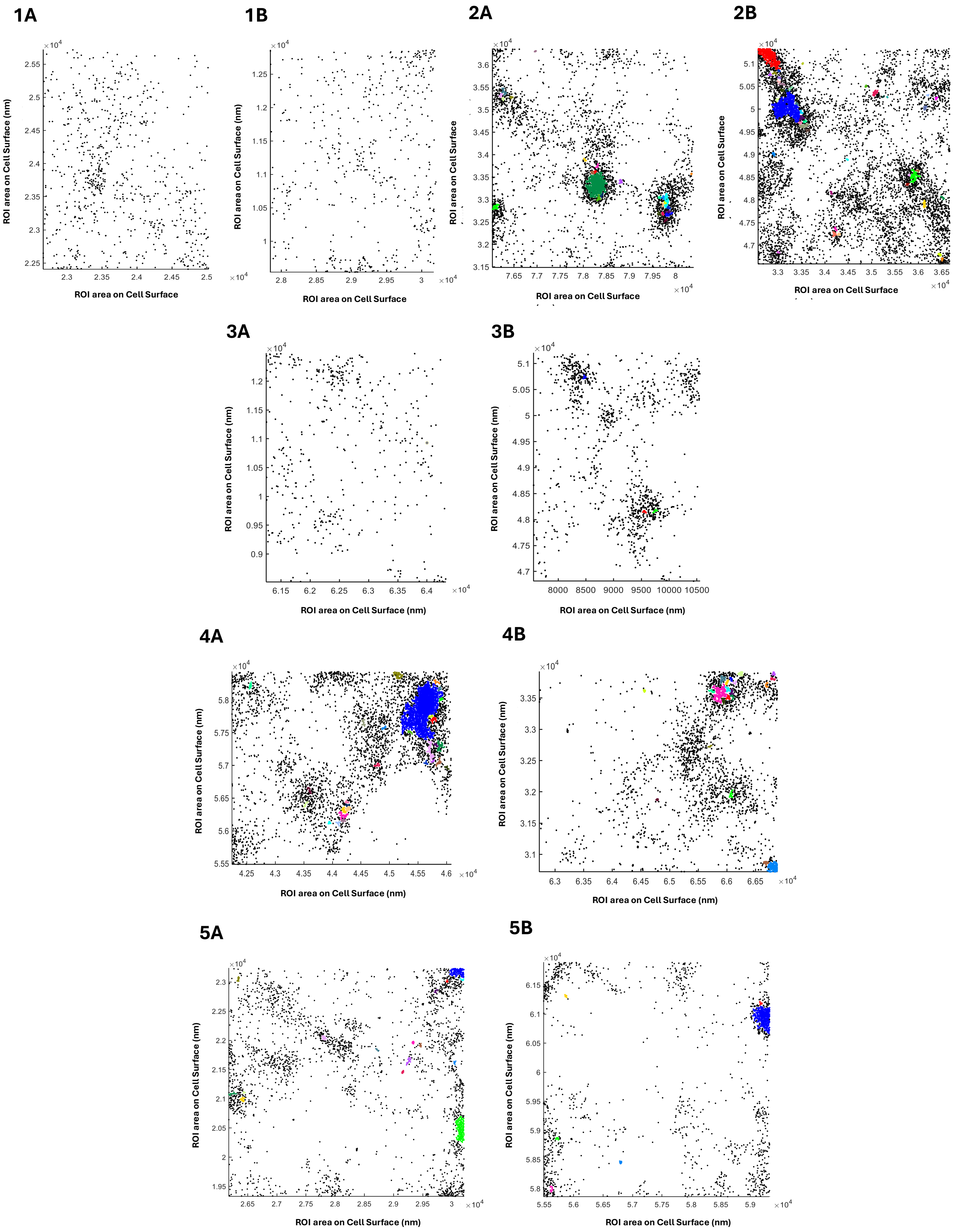

1A
ROI area on Cell Surface (nm)
ROI area on Cell Surface
1B
ROI area on Cell Surface (nm)
ROI area on Cell Surface
2A
ROI area on Cell Surface
ROI area on Cell Surface
2B
ROI area on Cell Surface
ROI area on Cell Surface
3A
ROI area on Cell Surface (nm)
ROI area on Cell Surface (nm)
3B
ROI area on Cell Surface (nm)
ROI area on Cell Surface (nm)
4A
ROI area on Cell Surface (nm)
ROI area on Cell Surface (nm)
4B
ROI area on Cell Surface (nm)
ROI area on Cell Surface (nm)
5A
ROI area on Cell Surface (nm)
ROI area on Cell Surface (nm)
5B
ROI area on Cell Surface (nm)
ROI area on Cell Surface (nm)

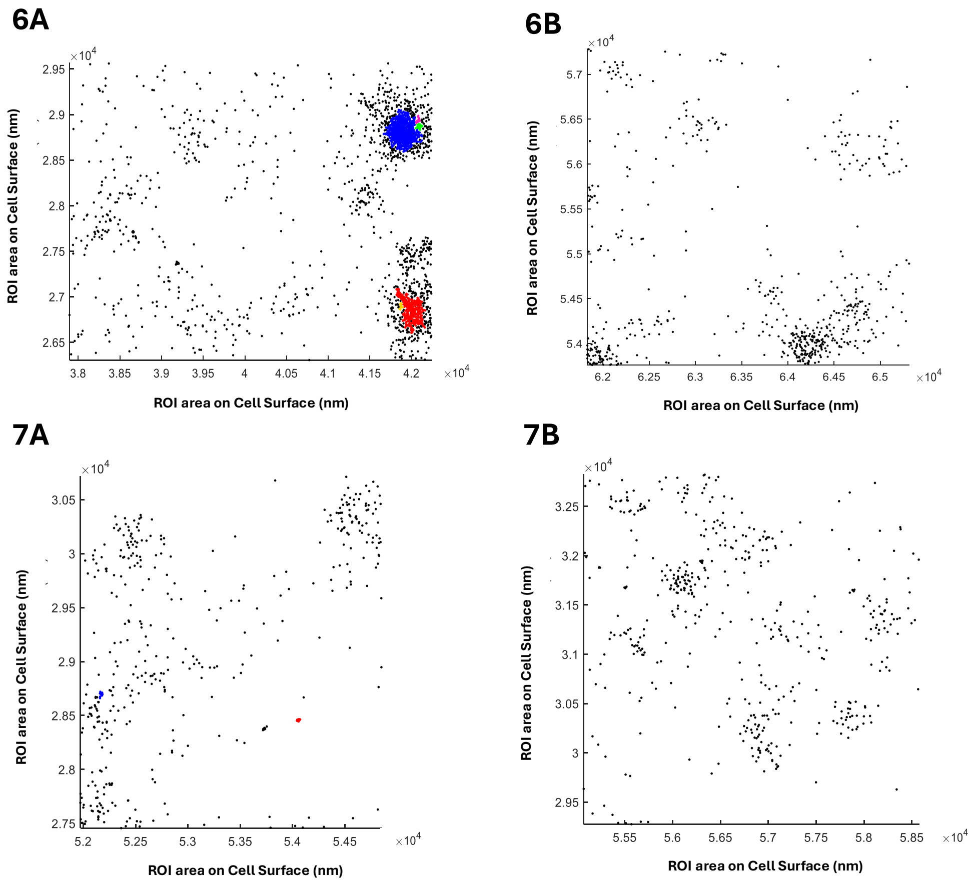


**Supplementary figure 8.** Surface TCR clusters were measured as per figure 6 and supplementary figure 7, with TCR cluster maps being produced in MATLAB post image acquisition. Images show duplicate cluster maps from 2 regions of interest (A and B) per conditions. Conditions shown: 1) Unstimulated 2) Immobilised CD3/CD28 control 3) No MF 4) 5min MF 5) 15min MF 6) 30min MF 7) 60min MF + 2hr wait. Images representative from 2 independent experiments. Colours shown on cluster maps indicate the positions of clusters identified by the DBSCAN software. Black points in the cluster maps indicate positions of TCR detections not allocated into a cluster. Cluster maps show formation of clusters in response to magnetic force application at 5 and 15 minutes, which then returns to baseline at 30 minutes and beyond.

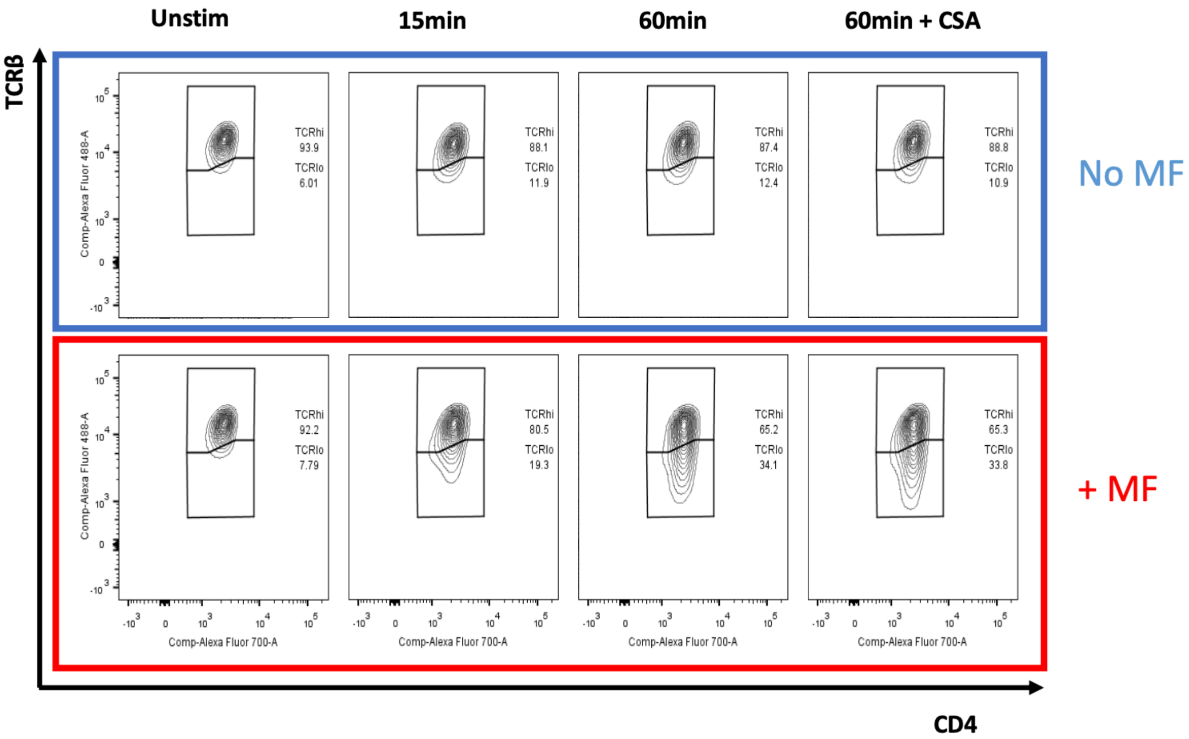


**Supplementary figure 9.** Blockade of the NFAT pathway with CsA fails to block TCR downregulation in response to 1 hour of magnetic force application. $CD4^+$ T cells were treated with 1μm anti-CD3 MNPs and subjected to magnetic force for the time frames shown. Where indicated, cells were also treated with 1μM CsA. Treatment with CsA failed to block force induced TCR downregulation.
